# Carvacrol as a dual-target acaricidal candidate against the cattle tick *Rhipicephalus (Boophilus) microplus*: molecular docking, mechanistic validation, and early field testing

**DOI:** 10.64898/2026.05.26.727743

**Authors:** Gloria S. Avendaño-Mora, Andrés F. Cabezas-Cárdenas, Bethsy N. Alfonso-Núñez, Erika M. Celis Celis, Cristhian Camilo Otero, Julián M. Botero Londoño, Wendy N. Triana, María Carolina Velásquez-Martínez, Stelia C. Mendez-Sanchez, Jonny E Duque

## Abstract

The blood-sucking ectoparasite *Rhipicephalus B. microplus* represents one of the greatest economic threats to livestock worldwide. The discovery of novel acaricidal molecules based on environmentally sustainable technologies is therefore required, particularly those suitable for use in dairy and meat-producing livestock without generating contaminant residues. This study aimed to identify and validate plant-derived metabolites with dual activity on acetylcholinesterase and mitochondrial targets as sustainable acaricidal agents against *R. microplus*, integrating computational screening, biochemical assays and field evaluation. Following *in silico* screening of 1.300 plant metabolites targeting acetylcholinesterase and mitochondrial function, fourteen compounds were selected for *in vitro* and *in vivo* mortality assays using immersion and larval vessel methods. Carvacrol was identified as the most promising metabolite and showed an inhibitory mechanism of action on acetylcholinesterase and mitochondrial complexes I, III, and IV. Its acaricidal efficacy was subsequently confirmed under field conditions using ethanolic and oily formulations (spray and pour-on) applied to cattle (*Bos taurus*). The spray formulation significantly reduced tick infestation, decreasing tick counts by 20% every 10 days (p < 0.001), whereas the pour-on formulation showed no significant reduction (p = 0.093). These results demonstrate that an integrated discovery pipeline from computational screening to field validation provides a robust strategy for identifying plant-derived acaricidal agents targeting key physiological pathways in *R. microplus*.

**Highlights:**

- Structured and sequential workflow (pipeline) from *in silico* to *Bos taurus*
- Identification and validation of phytometabolites against *R. microplus*
- Carvacrol multitarget activity on acetylcholinesterase and bioenergetic in ticks
- The spray formulation reduces tick infestation by 20% every 10 days

**Graphical abstract:** 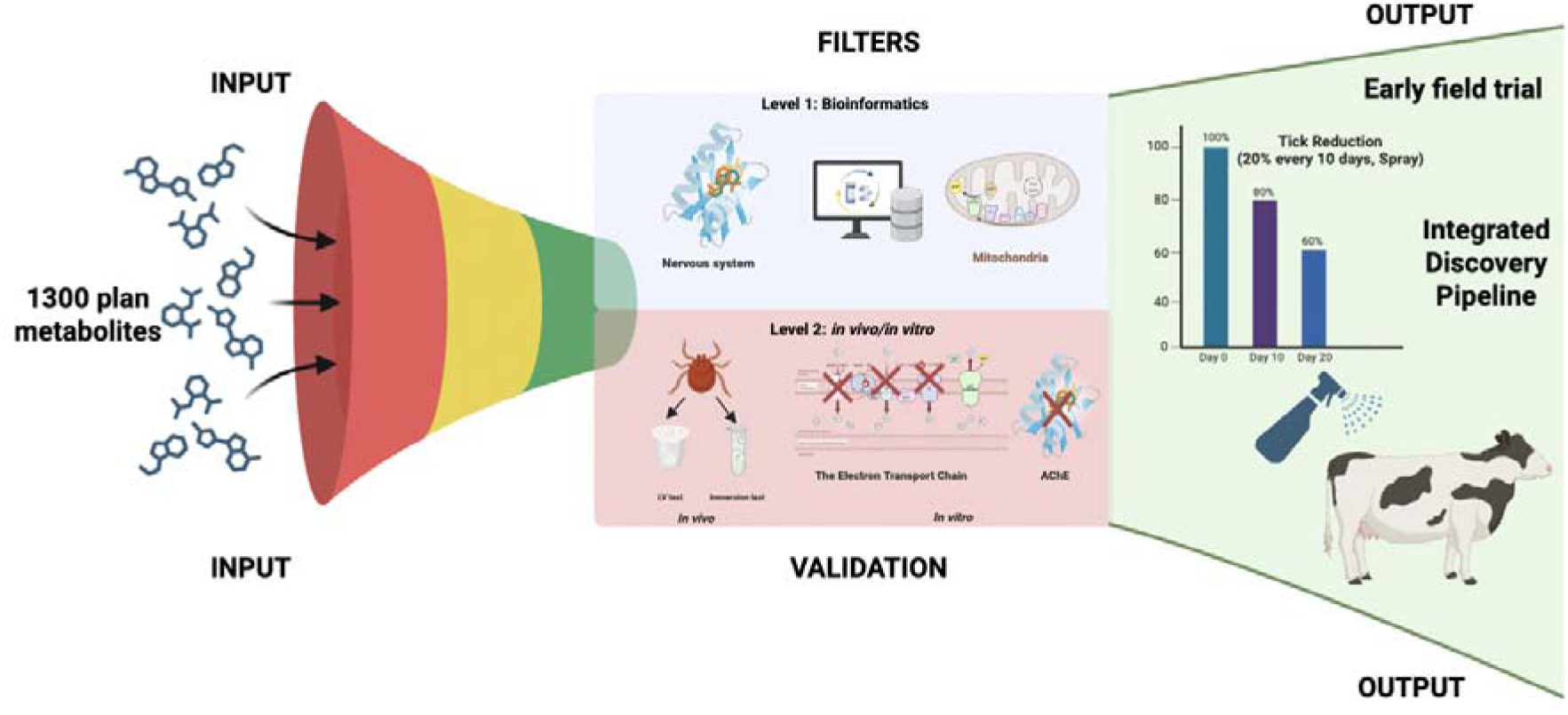

Created in BioRender. Avendaño, G. (2026) https://BioRender.com/7pm1ffj

## 1. Introduction

Ticks are blood-feeding parasites found in a wide range of climates, from arid regions to highly humid environments. They have a great capacity for survival and can withstand prolonged periods, even years, without feeding while waiting to find a suitable host (Boulanger et al., 2019). Their hosts can range from birds and reptiles to mammals, demonstrating their adaptability and ecological versatility (Rodríguez-Vivas et al., 2014). The tick species *Rhipicephalus microplus*, which is native to Asia, has spread to tropical and subtropical regions worldwide. This species has established itself as an invasive species of global concern due to its rapid, ongoing geographic expansion, with recent detections in Kenya and West Africa. This phenomenon is not only displacing native species such as R. decoloratus but also exacerbating risks to animal health and global livestock production (Kanduma et al., 2020; Makenov et al., 2021). By infesting approximately 80% of the world’s cattle, *R. microplus* is recognized as an effective vector of key blood parasites, such as *Babesia* spp. and *Anaplasma* spp., which cause serious diseases, including babesiosis and anaplasmosis (Polanco & Ríos, 2016). The epidemiological implications of the rapid spread of *R. microplus* are key to understanding the pathological and productive effects that this species causes in cattle. In this regard, infestations can cause allergic skin disorders, as well as weight loss, anemia, and reduced milk and meat production, which in severe cases can lead to the death of the animal (Adenubi et al., 2016). In fact, several studies have indicated that even a single engorged adult tick can reduce daily weight gain by 0.6 g and decrease milk production by approximately 8.9 mL per cow (Sutherst et al., 1983). Therefore, the presence of 20 or more adult ticks on a single animal represents a considerable risk to livestock health and has negative economic consequences for livestock production (Sepúlveda et al., 2017).

There are currently several methods for tick control, including traditional approaches such as synthetic acaricides, vaccines, and entomopathogenic fungi (Weber & Selzer, 2016). Additional management practices seek to interfere with tick biology and host–parasite interactions, including pasture rotation and the use of plant species with natural repellent properties. Essential oils and plant extracts have been proposed as alternative acaricidal agents; however, their efficacy in tick control under field conditions is often modest and inconsistent (Jaramillo H. et al., 2019; Malak et al., 2024). Consequently, chemical control remains the most widely adopted strategy, primarily relying on acaricides based on organophosphates, synthetic pyrethroids, and macrocyclic lactones. Despite their widespread use, these compounds present several limitations. Their efficacy often declines over time, leading livestock producers to increase dosages, apply treatments more frequently, rotate products without proper guidance, or combine different acaricides. Such practices accelerate the selection of resistant tick populations. In addition, some acaricides pose significant risks to both human health and the environment. Organophosphates, for instance, are among the leading causes of acute pesticide poisoning in humans (Robb et al., 2026), whereas synthetic acaricides may bioaccumulate in animal tissues, resulting in contamination of bovine-derived products such as milk and meat (Bravo-Guerrero et al., 2024; Yaima Yate & Díaz Rivera, 2022). Despite these concerns, the discovery of new acaricidal molecules remains slow and has not kept pace with the global expansion of tick infestations.

In this context, there is an urgent need to develop new, effective, and ecologically sustainable strategies for the control of *R. microplus* that are safe for human and animal health. Although less refined plant-derived products, such as crude extracts, have frequently shown limited and variable efficacy, growing evidence indicates that targeted exploration of plant secondary metabolites may overcome these limitations. Secondary metabolites isolated or enriched from complex botanical matrices represent a promising source of bioactive molecules with demonstrated activity against a wide range of pests and pathogens, offering structurally diverse scaffolds and multiple mechanisms of action that could counteract tick resistance to conventional chemicals (Yadav et al., 2022). Using secondary plant metabolites as ixodicidal agents has several advantages over synthetic acaricides (Malak et al., 2023). First, many of these compounds are readily biodegradable and exhibit lower environmental persistence, thereby reducing contamination risks and adverse effects on biodiversity (Shanmuganath et al., 2021). Second, their multitarget modes of action may reduce the likelihood of resistance development and prolong treatment efficacy (Celis Celis et al., 2026). Additionally, numerous plant-derived metabolites are supported by a long history of traditional use in medicine and agriculture, providing an important foundation of empirical knowledge that may facilitate their development, adoption, and acceptance as sustainable tick control agents (Adenubi et al., 2016; Quadros et al., 2020).

Recent efforts in pesticide discovery increasingly focus on identifying specific pharmacological targets that can be selectively modulated by natural compounds to generate safer and more effective bioinsecticides. In particular, enzymes involved in neurotransmission, such as acetylcholinesterase, and proteins associated with mitochondrial bioenergetics have emerged as promising molecular targets for the development of next-generation pesticides (Duque et al., 2023; Vanegas-Estévez et al., 2024). In mosquito vectors such as *Aedes aegypti*, studies integrating computational screening, biochemical assays, and biological validation have demonstrated that plant secondary metabolites can interfere with mitochondrial electron transport and acetylcholinesterase activity, leading to insect mortality (Castillo-Morales et al., 2021). Moreover, advances in experimental methodologies to evaluate mitochondrial respiration and enzymatic activity have provided valuable tools to elucidate the effects of xenobiotics on bioenergetic processes and to support the mechanistic characterization of candidate bioinsecticides (Vesga et al., 2025). These developments have established an integrated discovery framework in which computational prediction, biochemical validation, and biological assays are combined to identify new compounds targeting key physiological pathways. Within this context, extending this strategy to other arthropod pests such as *R. microplus* represents a promising approach for discovering novel acaricidal agents with defined mechanisms of action.

## 2. Materials and methods

### 2.1 Chemical Reagents

All reagents were of analytical grade, and solutions were prepared in Milli-Q water. The secondary metabolites eugenol, carvacrol, citral, gallic acid, naringenin, and quercetin, flavone, hesperidin, hydrobenzoin, 3-hydroxyflavone, syringaldehyde, trans-ferulic acid as well as 5,5’-Dithiobis(2-nitrobenzoic Acid) (DTNB), acetylcholine iodide, Monosodium phosphate (NaHLJPOLJ), succinic acid, 1,4-Dihydronicotinamide adenine dinucleotide (NADH), Ethylenebis(oxyethylenenitrilo)tetraacetic acid (EGTA), Ethylenediaminetetraacetic acid (EDTA), 2,6-Dichlorophenolindophenol (DCPIP), Phenazine methosulfate (PMS), potassium ferricyanide, rotenone, sucrose, 4-(2-Hydroxyethyl)-1-piperazineethanesulfonic acid (HEPES), Bovine Serum Albumin (BSA), and cytochrome c, were purchased from Sigma-Aldrich. Other chemicals, such as linalool, acetone, ethanol, potassium dihydrogen phosphate (KH_2_PO_4_), dipotassium hydrogen phosphate (K_2_HPO_4_), ammonium heptamolybdate ((NH_4_)_6_Mo_7_O_24_), and ferrous sulfate (FeSO_4_), were purchased from Merck. Mineral oil was obtained from Laboratorios León S.A.

### 2.2 *In silico* analysis

A list of 1,300 secondary metabolites characterized from plants with reported acaricidal, repellent, or insecticidal activity was initially compiled for molecular docking screening. In parallel, enzymes characteristic of each mitochondrial electron transport chain complexes and acetylcholinesterase were selected as drug targets. The steps for performing molecular docking are described below.

#### 2.2.1 Obtaining homology models

To date, the three-dimensional structures of the enzyme acetylcholinesterase and the mitochondrial protein subunits of *R. microplus* have not been characterized, so we performed homology modeling using AlphaFold software with the primary structure or FASTA sequence provided in UniProtKB30. The subunits selected for each mitochondrial complex and AChE were: NAD4 chain 4 of complex I (ID: V9MMD6), iron-sulfur protein of mitochondrial complex II (ID: A0A6M2CRB1), mitochondrial complex III Rieske protein (ID: A0A6M2CSX0), COX1 - complex IV (ID: V9MLX4) and AChE (ID: M1U2A0).

#### 2.2.2 Molecular docking

The ligands (metabolites) were prepared using Schrödinger, Inc.’s LigPrep tool. The ADME properties were obtained using the QikPRO tool. The proteins were prepared using the Protein Preparation Wizard tool from the same software. The possible interaction sites between the proteins of interest and the ligands were determined using the Sitemap tool in Maestro. Finally, the Grid Generation tool was used to define the specific area of the box within which protein-ligand docking was performed for each subunit; the coordinates of the box are provided in supplementary *Table S1*.

The Maestro Glide tool was used to perform high-throughput structure-based protein-ligand docking. The metabolite docking process was performed at two levels of accuracy. In the first stage, the standard precision mode (SP) was applied to the 1,300 metabolites that comprised the initial database. In this first phase, the best poses for each ligand were selected based on docking scores, enabling classification of 250 metabolites. In a second stage, extra-precision (XP) docking was performed, generating a larger number of conformations per ligand to enable a more comprehensive analysis of possible interactions with the proteins of interest. From this last level, the 100 compounds with the highest scores were selected. The protein-ligand affinity energy was determined for these compounds using the Prime MM-GBSA tool, optimizing calculated parameters and eliminating possible metal ions, hydrogen bonds, and solvent molecules. This analysis identified 53 compounds with the highest binding scores and affinities. Following, the ADME properties of this group were calculated, and based on compliance with Lipinski’s rules and low toxicity, only 14 metabolites (carvacrol, citral, eugenol, flavone, gallic acid, hesperidin, hydrobenzoin, 3-hydroxyflavone, linalool, naringenin, phthalic acid, quercetin, syringaldehyde, and trans-ferulic acid) were classified and advanced to the next stage of *experimental* validation.

### 2.3 *In vivo* experiments

Blood-engorged female ticks of the species *R. microplus* were collected from three different locations: the El Paraíso agro-ecotourism farm in the municipality of Betulia (coordinates 6°54′00″N 73°17′01″W), the La Llanada farm in the Junín district of the municipality of Concepción (coordinates 6°43′33.1″N 72°40′13.3″W), and the Jericó farm in the municipality of Floridablanca (coordinates 7°04′02.4″N 73°05′19.0″W). All these farms are located in the department of Santander, Colombia. The ticks were transferred to the medical entomology laboratory of the Center for Research on Tropical Diseases (CINTROP), where the species was identified taxonomically (Nava et al., 2019) and molecularly. The ticks were then cleaned and placed in Petri dishes lined with Whatman-type qualitative filter paper to facilitate oviposition. The eggs were collected in Eppendorf tubes lined with moistened cotton daily to maintain optimal humidity at 80-90%. The tubes were stored under a 12 h light/12 h dark photoperiod, at 25 ± 5°C and 70 ± 5% relative humidity.

For molecular identification, genomic DNA was extracted from tick samples using the DNeasy Blood & Tissue Kit (Qiagen, Germany), according to the manufacturer’s instructions. PCR amplification was carried out using three primer pairs targeting the mitochondrial COX-1 region (Table 1). Reactions were prepared with Q5 High-Fidelity 2X Master Mix (New England Biolabs, USA). PCR conditions included an initial denaturation step at 98°C for 30 s, followed by 35 cycles of denaturation at 98°C for 10 s, annealing at 67–71°C for 30 s using a gradient PCR approach to optimize conditions for each primer set, extension at 72°C for 30 s, and a final extension step at 72°C for 2 min. Amplicons of the expected size were excised from the gel and processed for downstream sequence analysis. PCR products were sequenced using Oxford Nanopore Technology (ONT). Libraries were prepared using the SQK-RBK110.96 Rapid Barcoding Kit, allowing unambiguous assignment of each amplicon through individual barcodes. Sequencing was performed on the PromethION P2 platform using an FLO-PRO114M flow cell, generating high-throughput sequencing data. Bioinformatic processing with wf-amplicon included raw-read refinement, adapter removal, alignment of each sequence set against its corresponding reference, and generation of Medaka-curated consensus sequences. Final inspection in Geneious confirmed that the obtained sequences corresponded to the target tick samples, with 100% identity in BLAST comparisons. Nucleotides used in this study were sourced from the project “Producción de nucleótidos a partir de biomasa residual de la agroindustria para diagnóstico por biología molecular en el departamento de Santander,” also approved by CEINCI under Minutes No. 20, dated May 20, 2022. Furthermore, the study was carried out in compliance with the arrive guidelines and following the 1989 Colombian Law 84 (Chapter IV, Art. 23–26) and Resolution 8430 (1993, Title IV, Art. 83–93) that regulates animal research in Colombia.

**Table 1.**
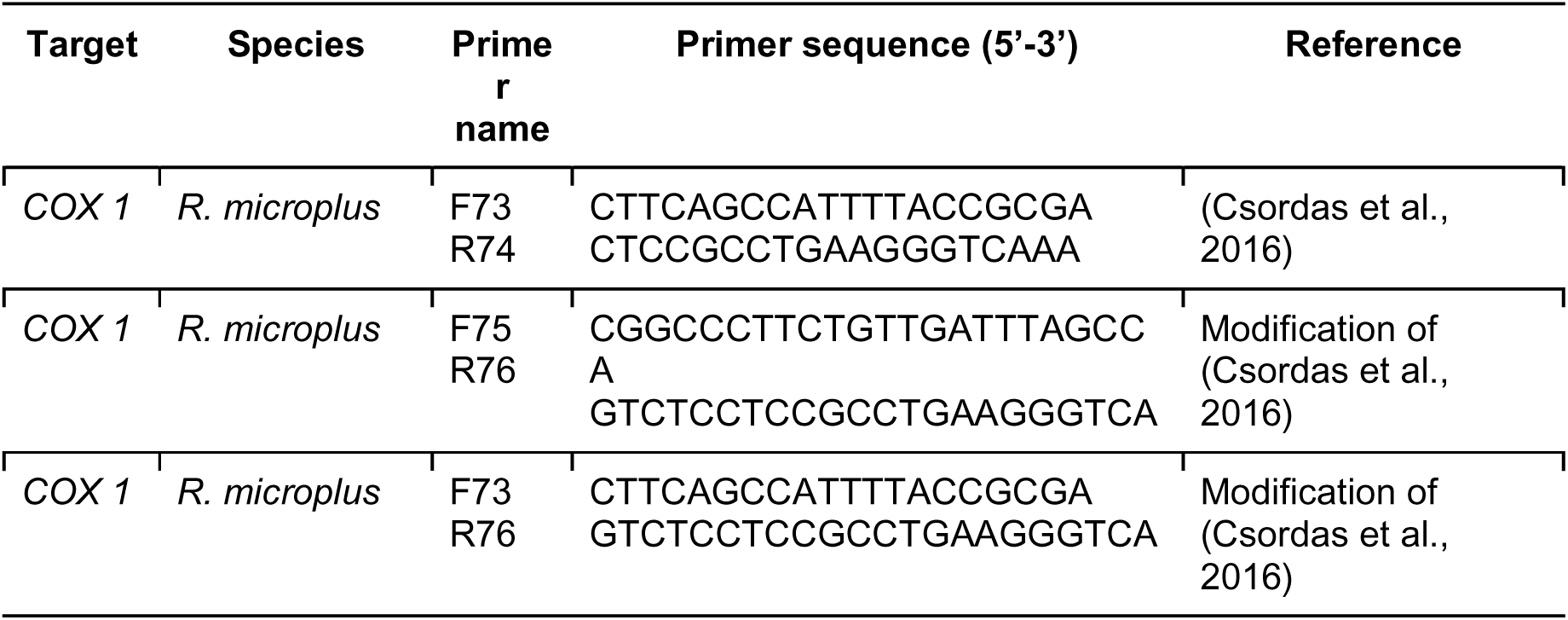
List of primers used for amplification of the *COX-1* gene.

#### 2.3.1 Acaricide testing using the larval vessel method

The mortality rate of *R. microplus* larvae between seven and fourteen days old was assessed across all assays. Four replicates were used for each concentration and target, evaluated over three different days. Dead individuals were counted at 24, 48, and 72 hours. A stereoscope was used to assess the mobility of the larvae; those that did not move within 2 minutes were counted as dead. Both the larval vessel and larval immersion methods underwent some modifications to adapt them to laboratory conditions. The first test was the larval vessel (LV) assay, an improved method reported for the first time in this study, based on the protocol described by (Adenubi et al., 2018), which was modified to extend the observation period for assessing the mortality of larvae exposed to the different test solutions. To do this, qualitative filter paper circles were used to construct a cone and a lid over a 1.5-ounce plastic cup. Each cone and lid was impregnated with 200 µL of solution, allowed to dry for two hours, and then 15 larvae were placed in each replicate (Figure 1). Three exploratory concentrations of each metabolite were initially evaluated: 100 ppm (0.66 mM), 365 ppm (2.43 mM), and 1000 ppm (6.66 mM). To obtain the lethal concentrations (LC_50_, LC_95_ and LC_99_), seven concentrations were evaluated for metabolites that showed an acaricidal response in the bioassay: 50, 100, 365, 500, 750, 1000, and 1300 ppm (equivalent to 0.33, 0.66, 2.43, 3.33, 5.00, 6.66, and 8.65 mM, respectively). The positive control was an ethion solution at 750 ppm (1.95 mM), and the negative control was acetone.

**Figure 1.**
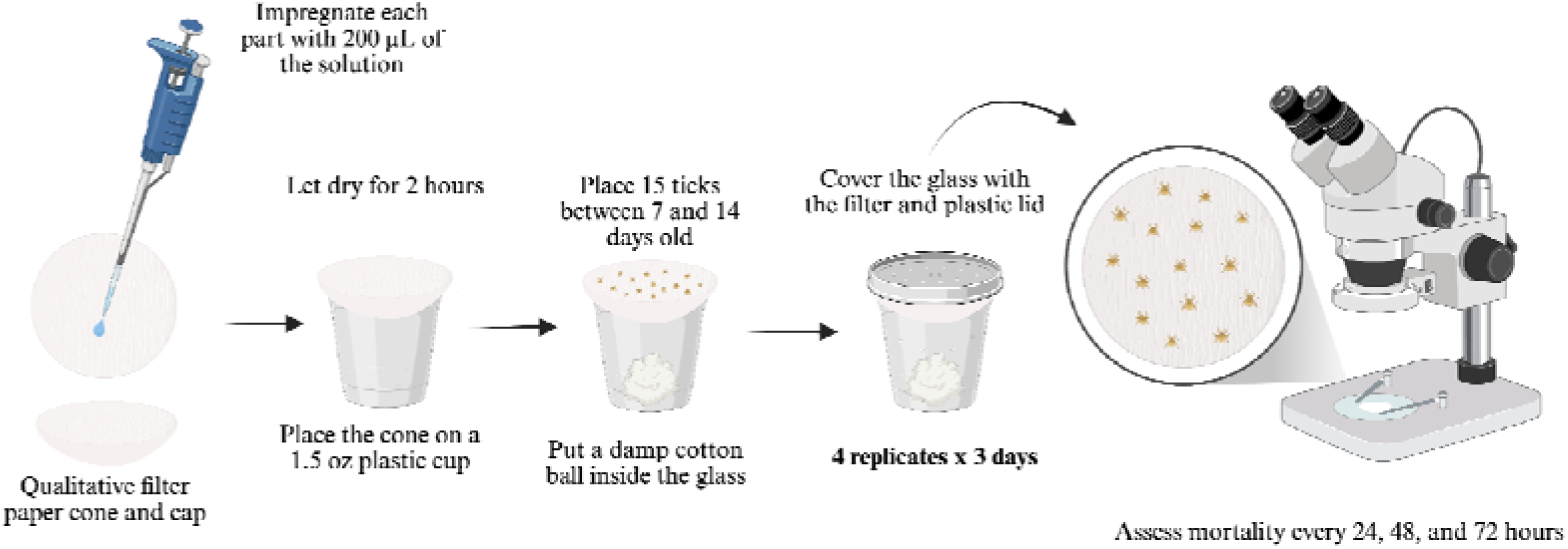
Experimental protocol to test acaricidal activity using the larval vessel. Created in BioRender. Avendaño, G. (2026) https://BioRender.com/8ykybun

#### 2.3.2 Acaricide tests using larval immersion

In this experiment, 25 larvae were used for each replicate. Four replicates were performed for each compound concentration. Live and dead larvae were counted at 24, 48, and 72 hours. Similarly, replicates were performed on three different days, for a total of 400 individuals analyzed. We followed the protocol of Klafke et al. (2021), modifying it to adapt to the conditions of the CINTROP laboratory, as shown in Figure 2 (Klafke et al., 2021). For the immersion tests, 10% ethanol was used as the solvent or negative control because acetone caused high mortality when larvae were immersed in it for 10 minutes (Novato et al., 2022; Ramírez et al., 2016; Sousa et al., 2022). Therefore, the 14 metabolites were evaluated using 10% ethanol in water as the solvent. Initially, three exploratory concentrations were evaluated for each metabolite (100, 365, and 1000 ppm). For the metabolite showing the highest mortality, seven additional concentrations (150, 320, 450, 550, 750, 900, and 1000 ppm corresponding to 0.99, 2.13, 2.99, 3.66, 4.99, 5.99, and 6.66 mM, respectively) were evaluated to determine lethal concentrations (LC values) comparable to those obtained in the LV assay using probit analysis.

**Figure 2.**
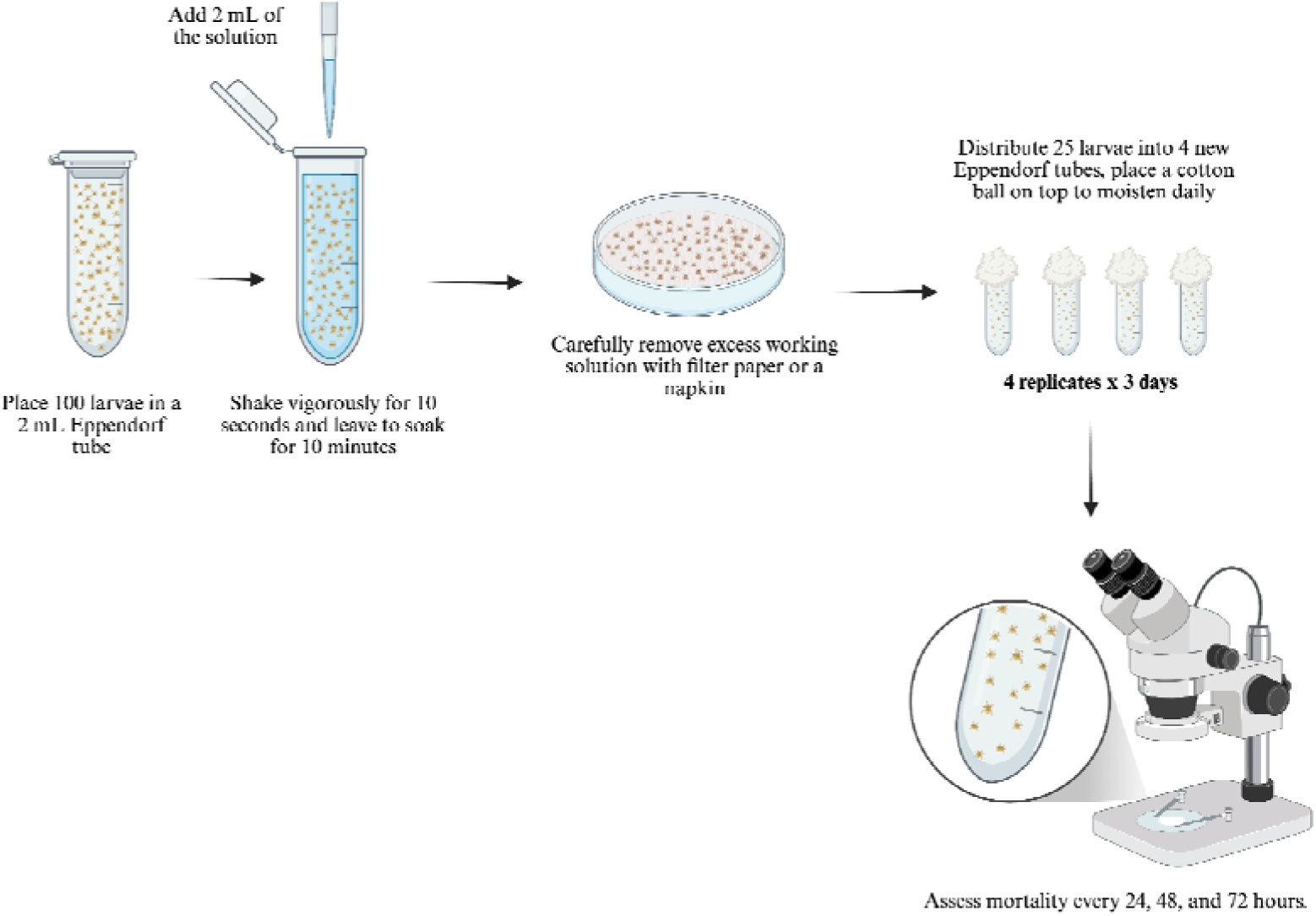
Modified experimental protocol for larval immersion acaricidal activity testing (Klafke et al., 2021). Created in BioRender. Avendaño, G. (2026) https://BioRender.com/8ykybun

To calculate the mortality percentage for each metabolite concentration, the following formula was used (Adenubi et al., 2018; Castro-Janer et al., 2009):

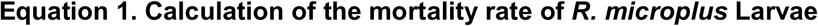

### 2.4 *In vitro* experiments

#### 2.4.1 Homogenization of *R. microplus* larvae

Approximately 500 mg of *R. microplus* larvae aged 21-60 days were refrigerated at 4°C for 10 minutes to anesthetize them. The larvae were placed in a porcelain mortar on an ice bed. Next, 1000 μL of homogenization medium, previosuly reported by Borrero et al. (2018) for *Aedes aegypti* mosquitoes, was added. This medium consisted of a solution containing 250 mM sucrose, 10 mM HEPES, and 1 mM EGTA, at pH 7.2 (Borrero Landazabal et al., 2018; Castillo-Morales et al., 2019; Duque et al., 2023). The larvae were manually macerated with a pestle until the chitin exoskeleton was completely broken down, yielding a viscous extract. The extract was then subjected to a series of centrifugations at 300 x g for 10 minutes to preserve the supernatant. Finally, total protein quantification was performed using the Bradford method (Bradford, 1976). For the initial NADH oxidase enzyme assays, a total protein concentration of 0.6 mg/mL was used, starting with a carvacrol concentration of 650 ppm (4.33 mM), a midpoint value located approximately between the LCLJLJ values obtained at 72 hours using the two acaricidal methods. Subsequently, the concentration was gradually reduced to determine the lowest level at which statistically significant differences were observed compared to the control. Based on these results, a concentration of 37.5 ppm (0.25 mM) carvacrol was selected for the subsequent assays.

### 2.4.2 Mechanism of action of carvacrol

The effect of carvacrol on mitochondrial electron transport chain enzymes and acetylcholinesterase was evaluated using a Multiskan GO spectrophotometer (Thermo Scientific, Waltham, MA, USA) with SkanIt software. In addition, the oxygen consumption of total proteins extracted from the larvae in the presence of the compound was quantified using a high-resolution Oroboros OLJk respirometer and DatLab CD software.

The effect of carvacrol on mitochondrial bioenergetic metabolism was evaluated through four experimental phases: 1) evaluation of inhibition of the entire electron transport chain through NADH and succinate oxidase assays and evaluation of inhibition of complexes I and II through NADH and succinate dehydrogenase assays using the Singer method (Borrero-Landazabal et al., 2020; Montagut et al., 2022; Singer, 2006); 2) evaluation of inhibition of complex III through NADH and succinate cytochrome c reductase assays using the Somlo method (Borrero-Landazabal et al., 2020; Somlo, 1965; Vanegas-Estévez et al., 2024) ; 3) evaluation of inhibition of complex IV through cytochrome c oxidase assays using the Mason method (Borrero-Landazabal et al., 2020; Mason et al., 1973) ; 4) evaluation of inhibition of ATP synthase using the Pullman method (Borrero-Landazabal et al., 2020; Pullman et al., 1960). The phosphate concentration was calculated using a calibration curve of phosphate standards at 1 and 300 μM, prepared before this assay (Borrero-Landazabal et al., 2020; Chen et al., 1956). In addition, 5) the effect of this metabolite on acetylcholinesterase, a second pharmacological target, was determined using the Ellman method (Castillo-Morales et al., 2019; Duque et al., 2023; Ellman et al., 1961).

### 2.5 Field pilot test in *Bos taurus* cattle using carvacrol solutions applied by spray and pour-on methodologies

#### 2.5.1 Field location and experimental conditions

The pilot field trial was conducted from July 23 to August 14, 2025, at the La Llanada farm (Coordinates 06°43’33.1” N; 72°40’13.3” W), located in the municipality of Concepción, Santander, Colombia, at an altitude of 1800-2800 meters above sea level. The area is characterized by a warm, mountainous, sub-paramo climate, with temperatures ranging from 8.5 to 19.5 °C and an average relative humidity of 80%.

### 2.5.2 Experimental Design and Field Procedures

A longitudinal field trial with repeated measures was conducted to evaluate the acaricidal efficacy of two carvacrol-based formulations under natural infestation conditions. The experimental unit was the individual animal, and each animal received a single treatment. Tick counts were recorded repeatedly over time, resulting in a completely randomized design with repeated measures, with treatment as a between-subject factor and time as a within-subject factor.

The study included twenty-four cattle of the *Bos taurus* (Creole) breed, ranging in age from 1 to 15 years; these animals were selected on a convenience basis. Prior to treatment, animals were maintained for up to 30 days without acaricide exposure to allow natural tick infestation. All individuals remained in their usual pasture under natural environmental conditions throughout the study. Baseline tick counts were performed on Day 0, with infestation levels ranging from 24 to 255 engorged ticks (> 3 mm in length) within a defined area on the left flank of each animal. Only engorged ticks larger than 3 mm were included to ensure standardized and biologically comparable measurements across sampling times. Animals were randomly allocated to four experimental groups (n = 6 per group): spray vehicle (0.5% v/v aqueous ethanol), pour-on vehicle (mineral oil), spray formulation (1500 ppm or 10 mM carvacrol in 0.5 % v/v ethanol), and pour-on formulation (1500 ppm or 10 mM carvacrol in mineral oil). Applications were conducted on two separate experimental days, with each treatment group subdivided into two replicates of three animals for operational feasibility. Following completion of the initial experimental phase, animals originally assigned to the spray vehicle group received the positive control treatment (ethion; commercial product Ganation®) at 1 mL per liter of working solution (final concentration 830 ppm or 2,16 mM), according to the manufacturer’s instructions.

Spray treatments were applied using a 20 L backpack sprayer, whereas the pour-on formulation was administered along the dorsal midline using a 50 mL syringe to ensure consistent topical delivery. Tick counts were recorded on Days 1, 2, 3, 4, 5, 7, 9, 11, 13, 15, 17, 19, and 21 post-application. At each time point, engorged ticks larger than 3 mm within the predefined anatomical area were counted. Animal health and welfare were monitored daily by a trained zootechnician through clinical assessment of body temperature, heart rate, body condition, lameness, behavior, and potential skin reactions at the application site.

On days 3 and 9 after the first application, ten (10) ticks were randomly collected from the spray control group, the spray formulation group, and the positive control. These ticks were transferred to the laboratory and stored in a Petri dish at 27 ± 1 °C and a relative humidity of ≥80%. Due to the heterogeneity and size of each group, only two parameters were observed as effects of the spray formulation: egg hatching rate and larval survival after hatching. To estimate hatching capacity, an aliquot of 50 eggs from each treatment unit was incubated in Eppendorf tubes. Equation 2 was used for the respective calculation.

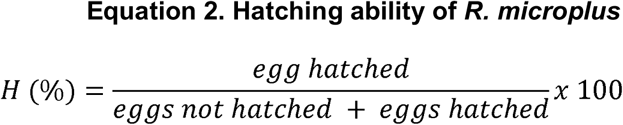

### 2.6 Statistical Analysis

Mortality data obtained from the metabolite bioassays were initially assessed for distributional assumptions using the Kolmogorov–Smirnov test (n > 50). Since the data significantly deviated from normality (p < 0.05), differences among treatments were analyzed using the Kruskal–Wallis test, followed by an appropriate multiple-comparison procedure for pairwise contrasts among treatments (Melo Martínez et al., 2020). Diagnostic concentrations of carvacrol were compared against both the negative and the positive control (Ethion). Statistical significance was set at p ≤ 0.05. For concentration–response analyses of carvacrol, mortality data from the LV and larval immersion bioassays were fitted to a Probit model using Polo Plus to estimate the median, 95%, and 99% lethal concentrations (LC_50_, LC_95_, and LC_99_), together with their corresponding 95% confidence intervals (Vanegas-Estévez et al., 2024). All analyses were conducted using Statistica 10.0.

Enzymatic activity data were expressed as mean ± standard deviation (SD) from three independent experiments, each performed with three replicates. Normality and homogeneity of variances were assessed prior to inferential analysis. When parametric assumptions were satisfied, differences among treatments were analyzed using analysis of variance (ANOVA), followed by Tukey’s post hoc test (Chong et al., 2018). Statistical significance was defined as p ≤ 0.05. These analyses were performed in Statistica 10.0 software.

In the pilot field trial, tick counts were analyzed using a negative binomial generalized linear mixed model (GLMM) with a log link to account for overdispersion. Treatment, time, and their interaction were included as fixed effects, with time scaled in 10-day units to facilitate biological interpretation of temporal changes. Sex and mean-centered age were incorporated as covariates. To account for repeated measurements within animals, individual animal identity was included as a random intercept. The significance of fixed effects was assessed using Wald z-tests. Treatment-specific temporal slopes were derived from the treatment × time interaction terms and expressed as percentage change over 10-day intervals using the transformation (exp(β) − 1) × 100. Pairwise comparisons among treatment-specific slopes were performed using Tukey-adjusted contrasts to control for multiple testing. Model adequacy was confirmed by evaluating dispersion, uniformity, and residual patterns. All analyses were conducted in R (version 4.4.1), with statistical significance set at α = 0.05.

The data on hatching and larval mortality obtained from ticks collected on day 10 from animals treated via spraying, as well as from the positive control group, were analyzed using a nonparametric framework. Prior to analysis, normality and homogeneity of variance assumptions were evaluated, and the data did not meet the criteria for parametric testing. Consequently, differences among treatments were evaluated using the Kruskal–Wallis test, followed by Dunn’s post hoc multiple-comparison test (Melo Martínez et al., 2020). Statistical significance was set at p < 0.05, and all analyses were conducted using Statistica.

### 2.7 Ethical aspects in the handling of ticks and cattle

This study was approved by the Scientific Research Ethics Committee (Comité de Ética en Investigación Científica, CEINCI) of the Universidad Industrial de Santander (Minutes No. 8 April 25, 2025). The handling of specimens and livestock complied with current Colombian regulations (Law 84/1989, Resolution 8430/1993, Decree 1843/1991), guaranteeing animal welfare, veterinary supervision, and controlled environmental conditions under the supervision of qualified professionals.

## 3. Results

### 3.1 Prediction of Secondary Metabolites with Acaricidal Activity in Acetylcholinesterase and Mitochondrial Electron Transport Chain proteins of *R. microplus* by Molecular Docking

Supplementary *Table S2* lists the metabolites in the database compiled from the plant species examined in the article (Benelli et al., 2016). This table correlates plant species, the country or region of study, the year, the mite species observed, the type of activity reported (repellent or acaricidal), and the identified metabolites. For some of the reported plants, secondary and tertiary searches were conducted to determine their chemical characterization in other sources; thus, additional articles were also included and correlated. The *in silico* process identified 53 potential metabolites that interact with different complexes of the electron transport chain and AChE (Supplementary *Table S3*). These metabolites belong to the following functional groups: polyphenols, flavonoids, terpenes, tannins, terpinenes, and allylbenzenes.

The metabolites selected for the *in vivo* assays are shown in Table 2. These metabolites were chosen for the next stage because they meet the Lipinski parameters provided by the Maestro software, are reported to be toxicologically safe in the literature, and are cost-effective to acquire.

**Table 2.** Selected metabolites for the *in vivo* acaricidal validation stage.

| PubChem<br>code | Molecule | Mol_MW<br><500 Da | DonorH<br>B <5.0 | AccpHB<br><10 | LogP<br>< 5.5 | Security | Cost<br>≤100<br>US\$ |
| --- | --- | --- | --- | --- | --- | --- | --- |
| 10364 | Carvacrol | 150.22 | 1.000 | 1.000 | 2.8 | Irritant | 144 |
| 638011 | Citral | 154.25 | 1.000 | 1.000 | 2.66 | Irritant | 90 |
| 3314 | Eugenol | 154.25 | 1.000 | 1.000 | 2.66 | Irritant | 48 |
| 10680 | Flavone | 222.243 | 0.000 | 2.500 | 3.00 | Irritant | 31.67 |
| 370 | Gallic acid | 170.12 | 4.000 | 5.000 | 0.21 | Irritant | 110 |
| 10621 | Hesperidin | 610.568 | 7.000 | 20.050 | 2.01 | - | 63.94 |
| 95447 | Hydrobenzoin | 214.26 | 2.000 | 3.400 | 1.70 | - | 93.59 |
| 11349 | 3-Hydroxyflavone | 238.242 | 1.000 | 3.250 | 3.40 | Irritant | 126.27 |
| 443158 | Linalool | 154.25 | 1.000 | 1.000 | 2.66 | - | 88 |
| 442428 | Naringenin | 272.25 | 3.000 | 5.000 | 1.84 | - | 47 |
| 1017 | Phthalic acid | 166.133 | 2.000 | 4.000 | 1.42 | Irritant | 37.35 |
| 5280343 | Quercetin | 302.24 | 5.000 | 7.000 | 1.23 | - | - |
| 8655 | Syringaldehyde | 182.176 | 1.000 | 4.250 | 1.10 | Irritant | 39.60 |
| 445858 | Trans-ferulic acid | 194.187 | 2.000 | 3.500 | 1.50 | Irritant | 47.51 |

**Note:** The abbreviations in these tables correspond to the following: Mol\_MW: Molecular Weight (130-500 Da), DonorHB: Number of Hydrogen bond Donors (0-5), AccpHB: Number of Hydrogen bond Acceptors (0-10), LogP: Octanol-Water partition coefficient ( $\leq 5.5$ ).

### 3.2. Molecular identification of tick specimens through COX-1 amplicon sequencing using Nanopore technology

Genomic DNA was successfully extracted from all tick specimens, yielding concentrations ranging from 483 to 531 ng/μL and A260/A280 ratios between 1.8 and 2.0, indicating high purity and suitability for downstream amplification. All three primer sets successfully amplified the COX-1 region across the analyzed samples. Gradient PCR identified 67°C and 71°C as the optimal annealing temperatures, depending on the primer set. Agarose gel electrophoresis confirmed amplicons of the expected size in all reactions, with no amplification observed in the negative controls (Figure S1). COX-1 amplicons from *R. microplus* samples S7374, S7576 and S7376 were sequenced on the PromethION P2 platform using an FLO-PRO114M flow cell. Sequencing produced 75.33k, 49.25k, and 91.39k reads, with passing read rates of 60.8%, 51.4%, and 57.1%, and total yields of 31.88 Mb, 23.86 Mb, and 36.51 Mb for S7374, S7576, and S7376, respectively (Table S4). The corresponding N50 values were 557 bp, 1,800 bp, and 523 bp. Bioinformatic processing with the wf-amplicon pipeline, including adapter trimming, per-barcode alignment, and Medaka-based consensus refinement, generated high-confidence consensus sequences for all samples. BLASTn analysis in Geneious Prime confirmed 100% identity with the target reference sequences, supporting the molecular identification of the tick specimens analyzed (Figures S2–S4).

### 3.3 Acaricidal Activity of 14 Promising Metabolites Against *R. microplus* Larvae

#### 3.3.1 Larval Vessel (LV)

Fourteen metabolites were initially screened at 100, 365, and 1000 ppm, resulting in mortality rates ranging from 0% to 66.7%. Thirteen compounds (citral, eugenol, flavone, gallic acid, hesperidin, hydrobenzoin, 3-hydroxyflavone, linalool, naringenin, phthalic acid, quercetin, syringaldehyde, and trans-ferulic acid) produced mortality below 20% after 72 h and did not differ significantly from the negative control at any concentration tested (Supplementary Table S5). These metabolites were therefore excluded from subsequent concentration–response assays.

Carvacrol was the only compound showing marked acaricidal activity, exceeding 50% mortality within 24 h at 1000 ppm and reaching 66.7% after 72 h. At 24 and 48 h, mortality at this concentration was significantly higher than that observed with ethion (Figure 3; Figure S5). A concentration and time-dependent response was observed for carvacrol (Figure 3). Mortality exceeded 50% at concentrations ≥750 ppm (4.99 mM) and reached 70–100% at 1000–1300 ppm (6.6–8.65 mM). Complete mortality was achieved at 1300 ppm after 48 h, exceeding the response obtained with ethion under the same conditions.

**Figure 3.**
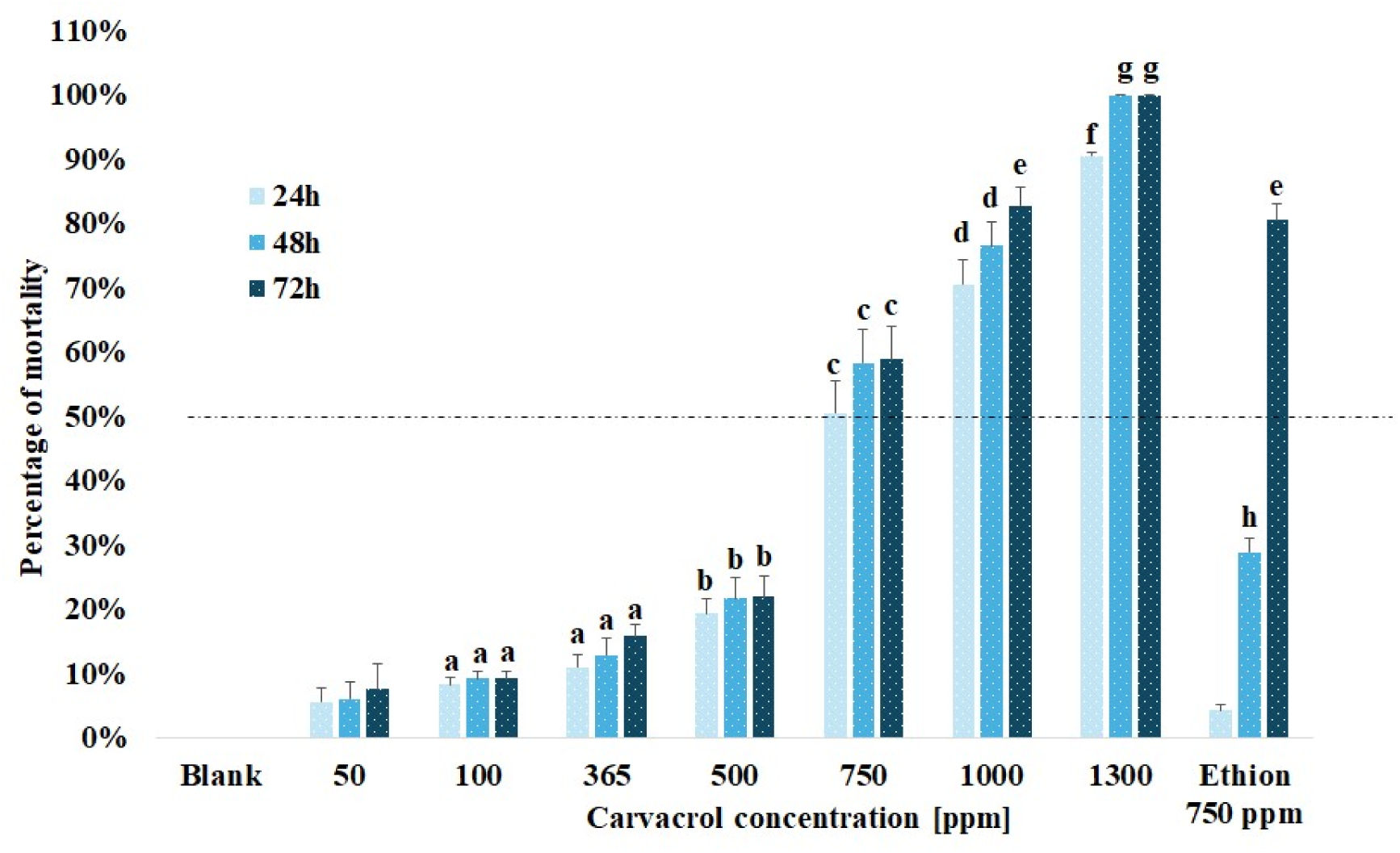
Mortality percentages for the LV test with carvacrol at seven concentrations are shown. Three independent quadruplicate trials were performed. Different letters above bars indicate significant differences among treatments (p ≤ 0.05).

Probit analysis yielded LC_₅₀_, LC_₉₅-_, and LC_₉₉_ values, which are presented in Table 3. Lethal concentration estimates decreased progressively from 24 to 72 h, with LC_₅₀_ declining from 755.5 to 682.6 ppm and more pronounced reductions observed for LC_₉₅_ and LC_₉₉_

**Table 3:**
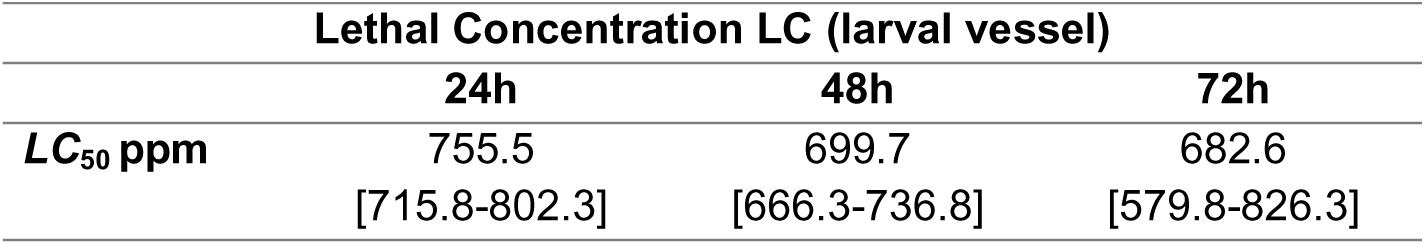
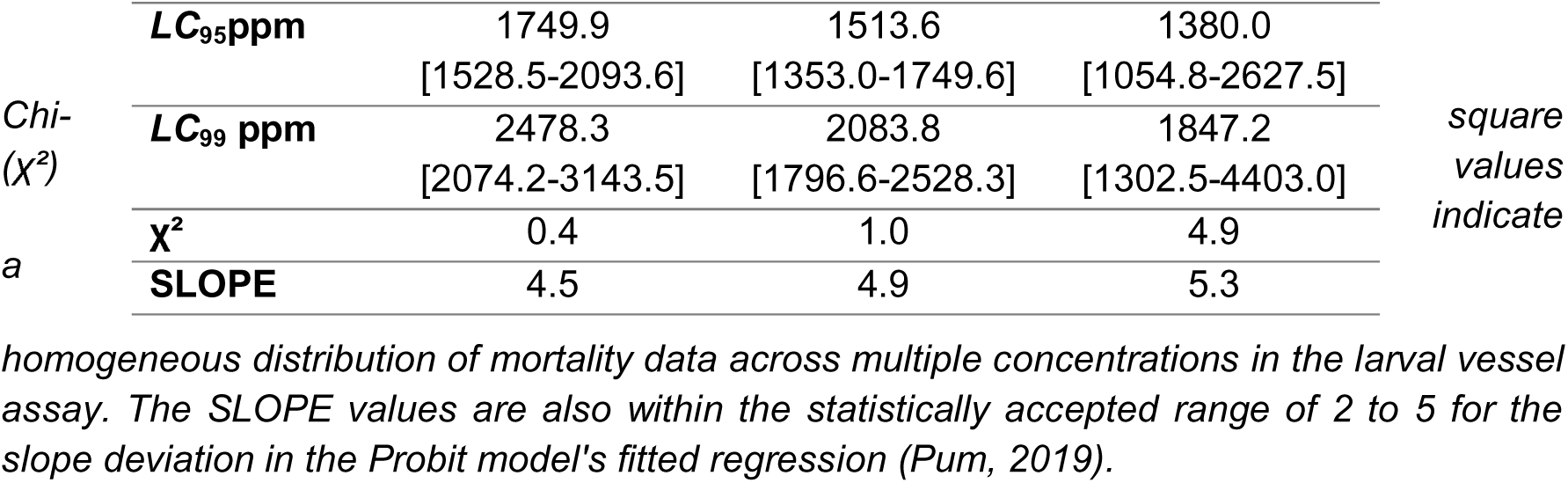
Data for lethal concentrations at 24, 48, and 72 hours for carvacrol per larval vessel (LV). Confidence intervals are shown in square brackets.

#### 3.3.2 Larval Immersion Test

Among the fourteen metabolites evaluated, only carvacrol showed relevant acaricidal activity in the larval immersion assay. The remaining 13 compounds did not exceed 10% mortality at 1000 ppm after 72 h and were excluded from further analysis (Supplementary Table S5). At the diagnostic concentration, carvacrol induced 100% mortality within 24 h, differing significantly from the negative control (Figure 4). Based on this response, concentration–response assays were performed at 150–1000 ppm (1.00–6.66 mM). Mortality increased progressively with both concentration and exposure time, similar to the larval cup method (Figure 4). At concentrations below 320 ppm (2.13 mM), mortality remained below 15%, suggesting an effective metabolic detoxification mechanism in the larvae, whereas approximately 50% mortality was observed at 750 ppm (4.99 mM) after 72 h. At 900 ppm (5.99 mM), mortality exceeded 80% within 24 h, and complete mortality was reached at 1000 ppm (6.66 mM), exceeding the response observed with ethion.

**Figure 4.**
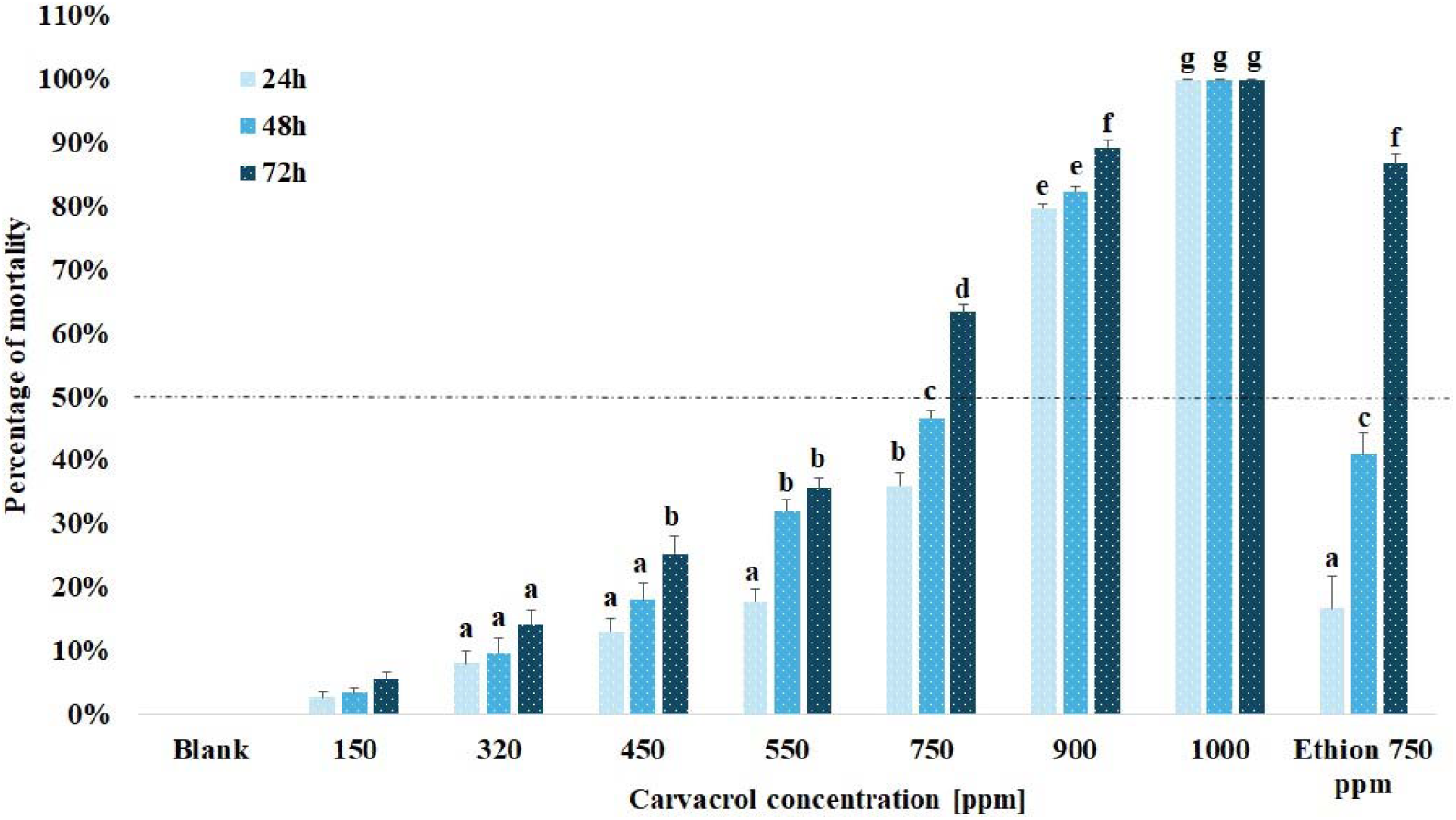
Mortality percent obtained in the larval immersion test with carvacrol + 10% ethanol at 7 concentrations is shown below. Three independent quadruplicate assays were performed. Different letters above bars indicate significant differences among treatments (p ≤ 0.05).

Probit analysis yielded LC_₅₀_, LC_₉₅_, and LC_₉₉_ values, which are summarized in Table 4. LC_₅₀_ values decreased from 717.9 ppm at 24 h to 570.0 ppm at 72 h. Compared with the LV assay, the immersion method consistently produced lower LC_50_ estimates across all exposure periods, suggesting increased larval susceptibility under immersion conditions. LC_₉₅_ and LC_₉₉_ values exhibited moderate variation across exposure times, remaining within comparable ranges.

**Tabla 4:**
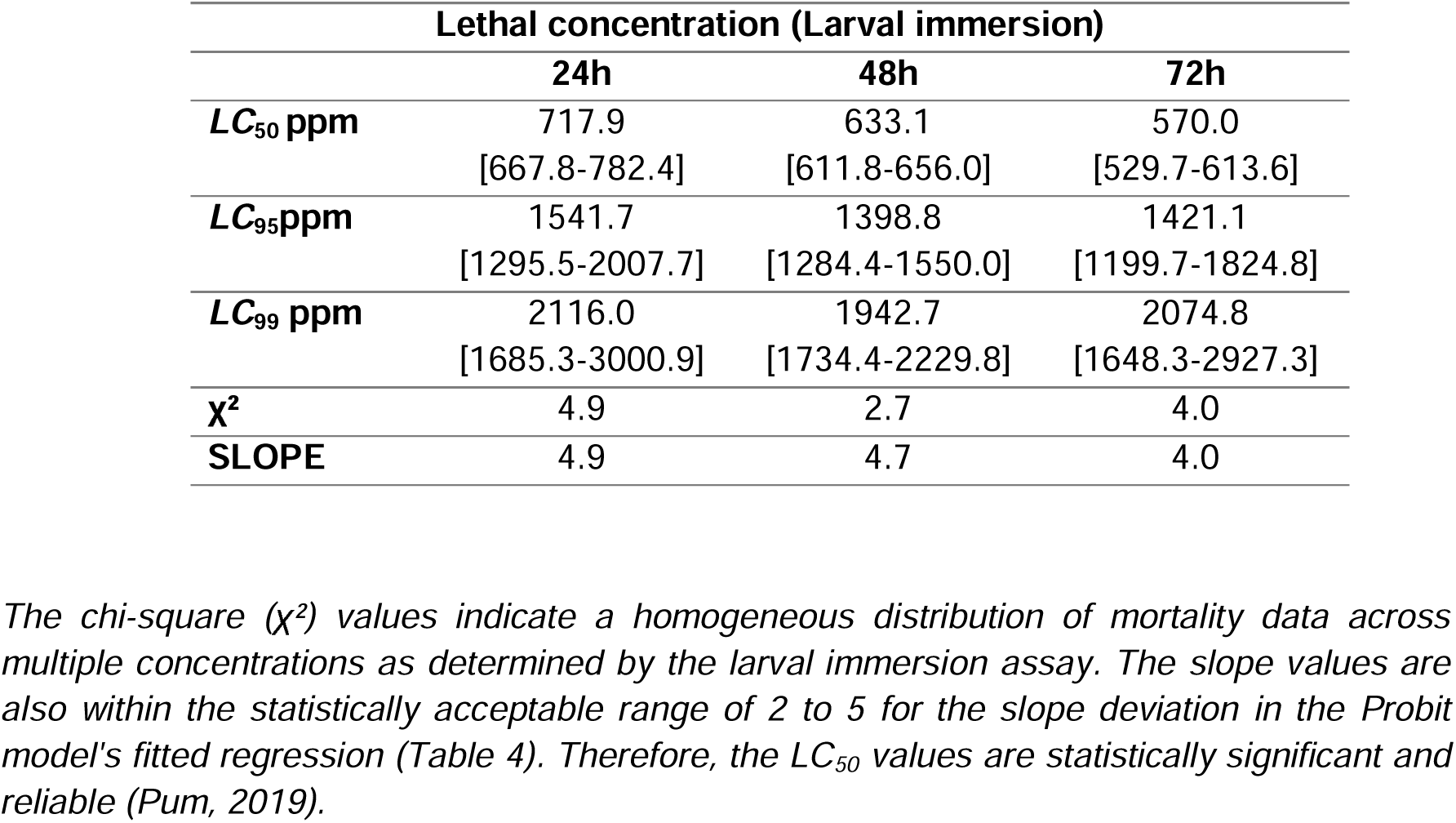
Lethal concentrations at 24, 48, and 72 hours for carvacrol by larval immersion assay. Confidence intervals are shown in square brackets.

### 3.4. Determination of the Mechanism of Action of Carvacrol on Mitochondrial of the Electron Transport Chain Proteins and Acetylcholinesterase of *R. Microplus*

#### 3.4.1 NADH Oxidase and Succinate Oxidase

Carvacrol inhibited NADH oxidase in a concentration-dependent manner (Figure 5). At 650 ppm (4.33 mM), corresponding to the average LC_₅₀_, activity was reduced by 76.2 ± 1.1%. The lowest concentration that produced a significant effect was 37.5 ppm (0.25 mM), which decreased activity by 11.9 ± 1.0%; this concentration was selected for subsequent assays. Control NADH oxidase activity was 31.54 ± 2.52 pmol/(s·mL). Succinate oxidase showed substantially lower activity than NADH oxidase under the same protein conditions (Figure S6), and the protein amount required to obtain measurable oxygen consumption exceeded the available extraction yield; therefore, this assay was not pursued further.

**Figure 5.**
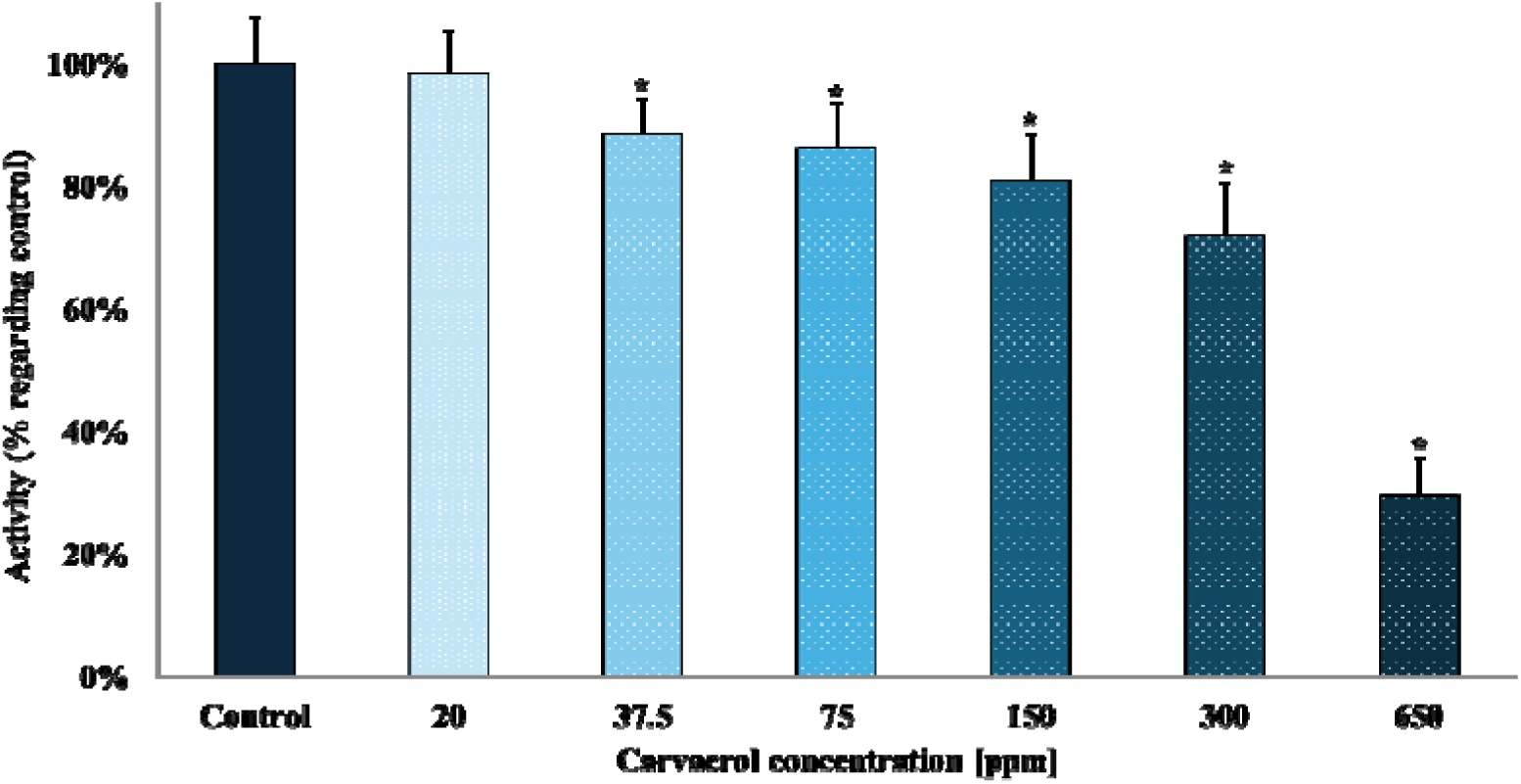
Effect of carvacrol on NADH oxidase activity. Enzyme activity was measured in the presence of increasing concentrations of carvacrol. The control value was 31.54 ± 2.52 pmol/(s·mL), corresponding to 100% activity. Values are expressed as mean ± SD. Asterisks (*) indicate significant differences relative to the control (p ≤ 0.05).

#### 3.4.2 Mitochondrial Enzymes and Acetylcholinesterase

At 37.5 ppm (0.25 mM), carvacrol reduced NADH dehydrogenase activity by 18.8 ± 1.7% relative to the control, indicating impaired electron transfer through complex I and decreased NADH oxidation. Succinate dehydrogenase activity remained unchanged, confirming that complex II was not affected (Figure 6). Cytochrome c reductase activity decreased by 30.0 ± 3.3% with NADH and by 23.1 ± 4.6% with succinate, identifying complex III as a major target of inhibition. Cytochrome c oxidase activity was also reduced, whereas ATP synthase activity showed no significant differences compared with the control, as confirmed by constant inorganic phosphate levels.

**Figure 6.**
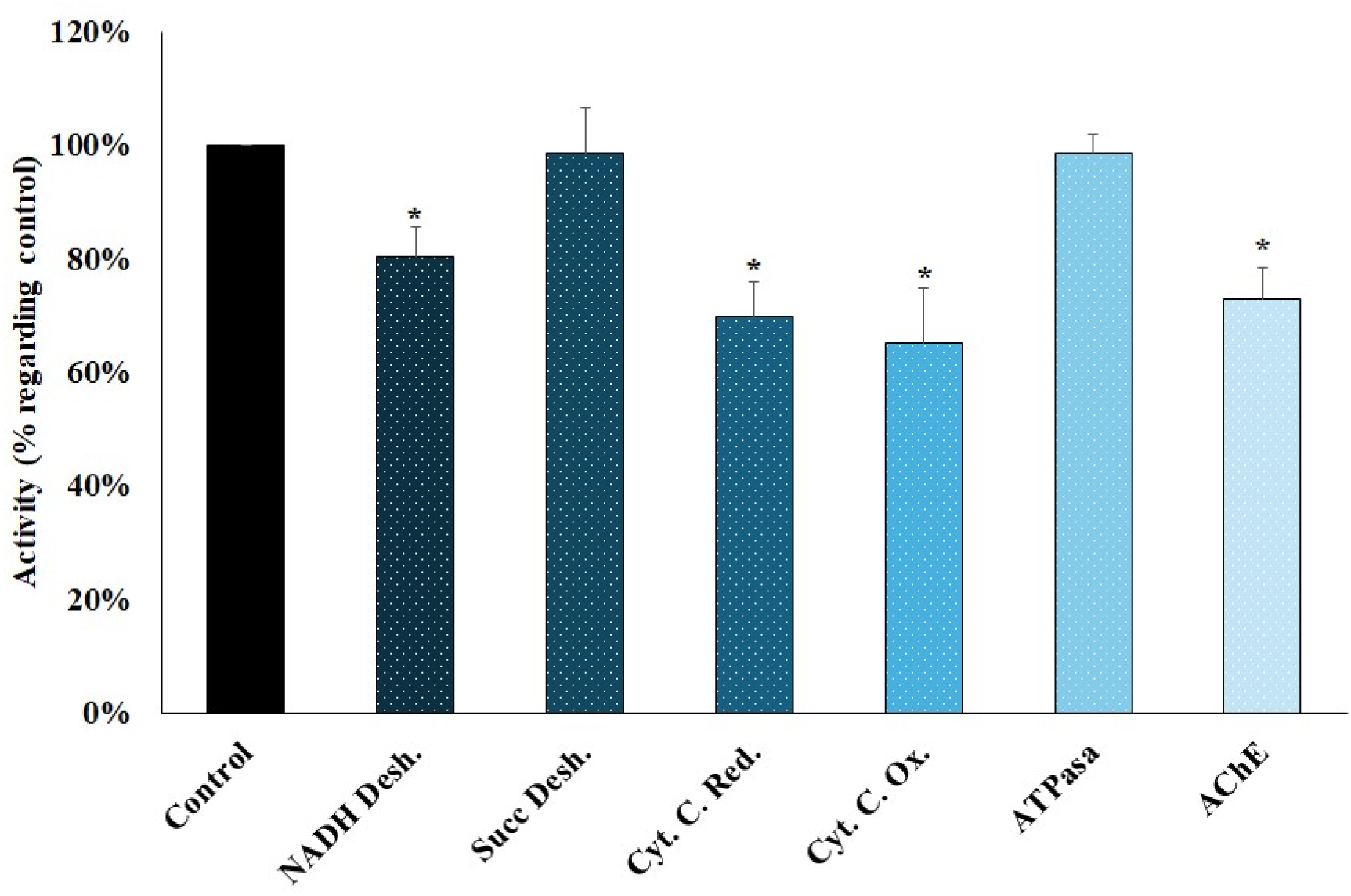
Effect of carvacrol at 37.5 ppm (0.25 mM) on mitochondrial electron transport chain enzymes and acetylcholinesterase activities. Enzyme activities are expressed as mean ± SD. The control was set at 100% and was found to be 556.6 ± 57.7 nmol of ferricyanide reduced/min*mg of tick protein for NADH dehydrogenase; 92.6 ± 3.9 nmol DCPIP reduced/min*mg tick protein for succinate dehydrogenase; 59.1 ± 4.68 nmol of reduced cytochrome C/min*mg tick protein for NADH cytochrome C reductase; 38.9 ± 4.2 nmol cytochrome c reduced/min*mg tick protein for succinate cytochrome C reductase; 40.6 ± 3.7 nmol cytochrome c reduced/min*mg tick protein for cytochrome oxidase; 0.58 ± 0.02 nmol of phosphate produced/mg tick protein for ATPase and 0.87 ± 0.07 nmol TNB produced/min*mg tick protein for acetylcholinesterase. Asterisks (*) indicate significant differences relative to the control (p ≤ 0.05).

Overall, carvacrol impaired electron transport chain efficiency, primarily affecting complexes I, III, and IV, while complex II and ATP synthase remained unaffected. In addition, carvacrol significantly inhibited acetylcholinesterase activity by 27.0 ± 2.9% relative to the control, suggesting potential neurotoxic effects through reduced acetylcholine hydrolysis (Figure 6).

### 3.5 Early field trial with carvacrol as an active ingredient in *Bos taurus* cattle

Tick infestation dynamics varied significantly among treatments over time (Figure 7). The negative binomial generalized linear mixed model (GLMM) identified a significant interaction between treatment and time (*Table S6*, Supplementary Material). In this case, the spray formulation showed a significant reduction in tick infestation, with a 20.0% decrease every 10 days (p < 0.001; *Table S6*). In contrast, the pour-on formulation did not show a significant reduction, with a decrease of only 5.0% every 10 days (p = 0.093; *Table S6*). Both negative control groups (5% ethanolic solution for spraying and mineral oil for pour-on) showed significant increases in tick numbers over time (Figure 7). Tick infestation increased by 15.1% in the spray control group (p < 0.001) and by 20.5% in the pour-on control group (p < 0.001; *Table S7*) every 10 days, respectively. The group treated with the positive control (ethion) also showed a significant increase of 12.0% every 10 days (p < 0.001).

**Figure 7.**
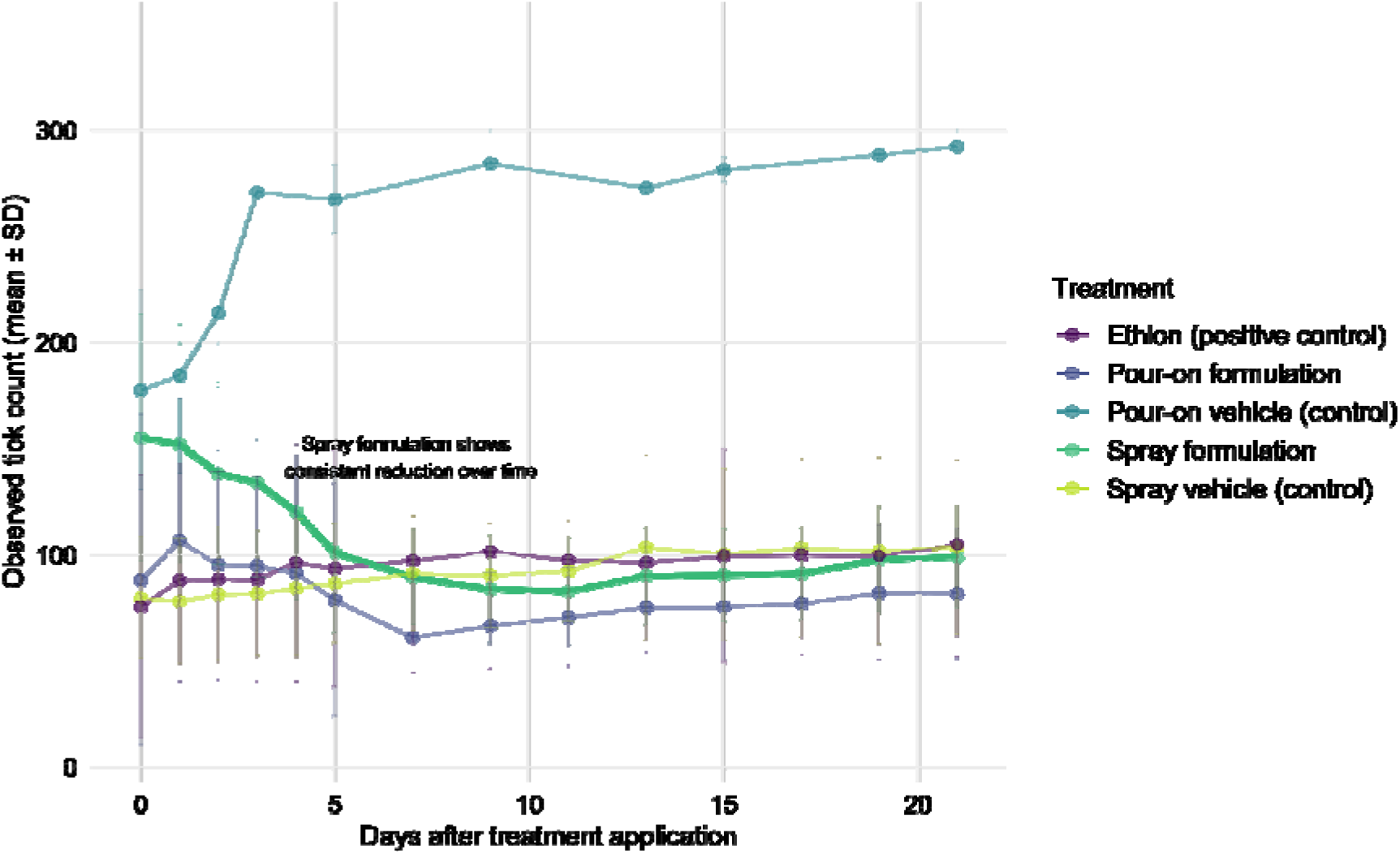
Observed tick infestation dynamics across treatments over time. Points represent mean tick counts at each sampling day, and error bars indicate ± standard deviation. Lines connect consecutive observations to illustrate temporal trends within each treatment group. The spray formulation exhibited a consistent reduction in tick counts over time, whereas other treatments showed stable or increasing infestation levels.

Pairwise comparisons of the time slopes (*Table S8* and *figure S7*, Supplementary Material) confirmed that the rate of change observed for the spray formulation differed significantly from that of all other treatment groups and the controls. No significant differences were detected between the pour-on formulation, the ethion-treated group, and the negative control group.

Regarding the monitoring parameters for engorged ticks treated with the active compound in spraying, the formulation achieved 96.6% inhibition of larval hatching (p < 0.001). In the negative control, inhibition was approximately 40%. Among hatched larvae, survival was only 0.4% following treatment (p < 0.0001), whereas ∼45% survived in the negative control (Figures 8A and 8B).

**Figure 8.**
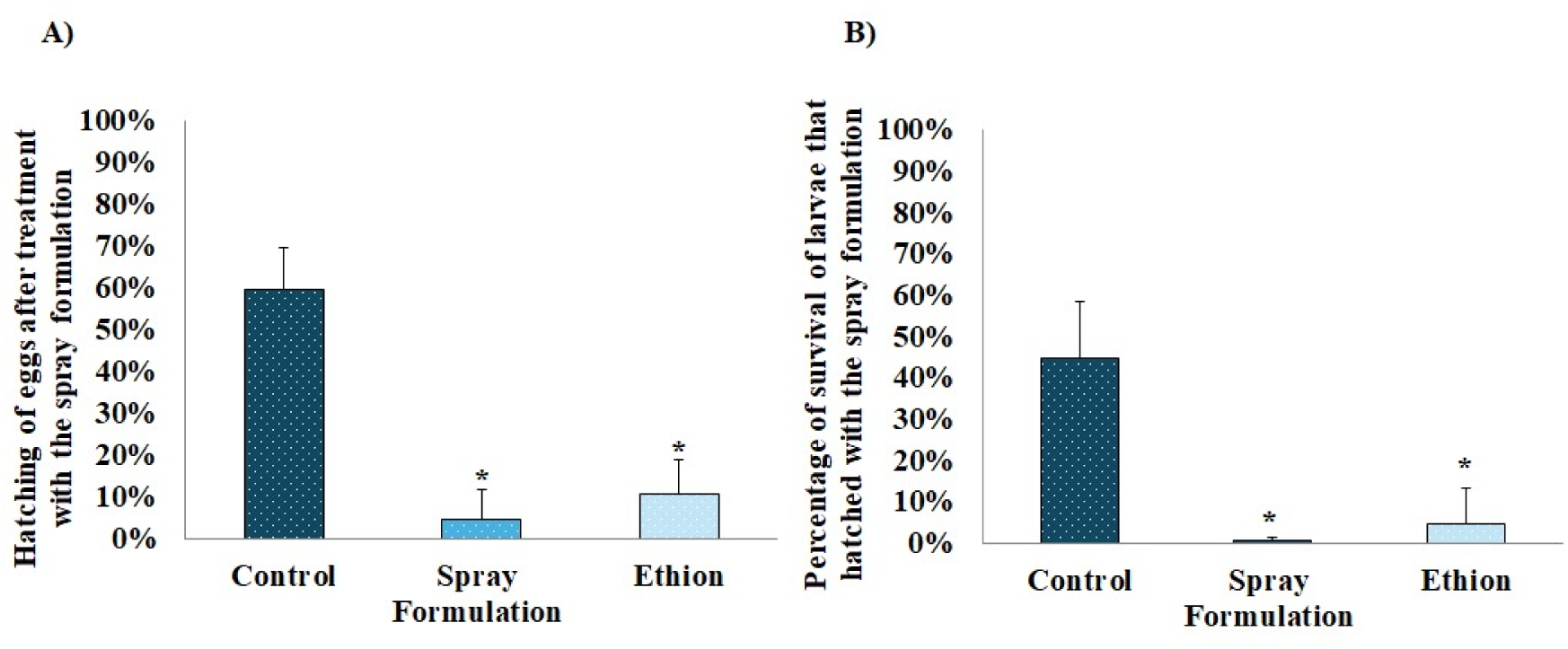
A) Hatching of eggs after treatment with the spray formulation. B) Percentage of survival of larvae that hatched with the spray formulation. Comparisons with positive and negative controls are shown, with significant differences marked with an asterisk (*).

## 4. Discussion

The discovery of molecules with acaricidal activity requires a structured methodological pipeline capable of integrating multiple criteria beyond direct toxicity against the target pest. In the current landscape of next-generation pesticides, mortality alone is no longer sufficient to support the development of a candidate molecule. Modern acaricidal products are expected to combine efficacy with selectivity, environmental compatibility, reduced persistence, and minimal risks to human, animal, and ecosystem health. Accordingly, this study implemented a stepwise discovery pipeline to identify plant-derived acaricidal candidates against *R. microplus*. This methodological strategy integrated in silico prioritization, in vitro validation, and an early field assessment, thereby enabling the evaluation of selected candidates beyond controlled laboratory conditions and providing preliminary evidence of their practical feasibility (Malak et al., 2024; Mendez-Sanchez et al., 2021; Vanegas-Estévez et al., 2024).

In this context, the initial library of 1,300 plant secondary metabolites was screened against two target classes that are directly linked to tick survival, enzymes of the mitochondrial electron transport chain and acetylcholinesterase (AChE), both previously reported as valid pharmacological targets for acaricide discovery. From the in silico screening, 53 metabolites showed potential interaction with one or more targets, and Lipinski-based filtering reduced the list to 14 compounds suitable for biological testing. Among these, carvacrol consistently demonstrated strong acaricidal activity in larval assays. This result corroborates earlier reports identifying carvacrol as one of the most active monoterpenes against *R. microplus*, with high mortality even at low concentrations, activity across resistant field populations, and enhanced effects in combination or formulation studies (Anjos et al., 2024; Cardoso et al., 2020).

It is worth noting that the current trend in the use of computational tools is aligned with research on acaricides focused on *R. microplus*, where platforms such as Schrödinger’s Maestro and AlphaFold stand out. In this context, the most recent study by Celis Celis et al. (2026) on NADH dehydrogenase subunit 4 (UniProt: V9MMD) is significant in supporting the findings of this study. The authors demonstrated hydrogen bond-like interactions between the hydroxyl group and the carbonyl oxygen of alpha-tocospiro A, and between the amino acids Ile247-Met250 and Gly249, respectively, of the mitochondrial enzyme from *R. microplus* (Celis Celis et al., 2026). In this study, carvacrol also showed affinity for the same complex I protein through polar interactions between the hydroxyl group of its structure and the amino acids Gly249, Ile247, and Met250, similar to that of alpha-tocospiro A. This molecular specificity could explain the observed alteration in the electron transport chain during mitochondrial respiration assays.

These computational models also support the identification of new acaricides that target the acetylcholinesterase (AChE) enzyme of *R. microplus* through electrostatic interactions and hydrogen bonds with residues in the active site, such as Ser202, Ser222, Ser303, and His460, for alkaloids such as juliprosinine and proserpine (Lima et al., 2020). This binding capacity has recently been further explored by Bustos et al. (2024), who, using AlphaFold structural models, identified metabolites in the seed extract of *Randia aculeata*, such as rutin and quercetin, that block this enzyme by interacting with His494, a specific residue of the catalytic triad (Bustos-Baena et al., 2024). It is worth noting that these *in silico* predictions were validated through in vivo assays on engorged adult *R. microplus* ticks, in which nervous system impairment was observed with tangible acaricidal efficacy. This established an important precedent for the transition from bioinformatics to its application in the field (Cerqueira et al., 2022; Concepción et al., 2013; Rodriguez-Vivas et al., 2018).

Acetylcholinesterase (AChE) is the second pharmacological target of carvacrol (Figure 9) in this study, with an enzymatic inhibition of 27.02 ± 2.95%. This inhibition is based on the architecture of the catalytic triad (Ser-His-Glu), which has been extensively characterized in insect acetylcholinesterases (Zhang et al., 2002; Zhou et al., 2010), and has also been identified in *Dermacentor variabilis*, with conservation of the catalytic residues across multiple tick species, including *R. microplus* (Cardoso et al., 2020). Carvacrol blocks the active site through a dual mechanism: the hydroxyl group forms a hydrogen bond with the Ser230, Gly150 and Gly151 residues, while its aromatic ring interacts with the imidazole ring of His476 via π-π interactions. This blocking not only physically restricts the access of the natural substrate to the catalytic center but also destabilizes the proton transfer necessary for acetylcholine hydrolysis and, consequently, disrupts synaptic communication in the tick. Therefore, the synergy between the impairment of mitochondrial bioenergetics and the disruption of synaptic communication explains the response to the spray-applied formulation in the early field trial.

**Figure 9.**
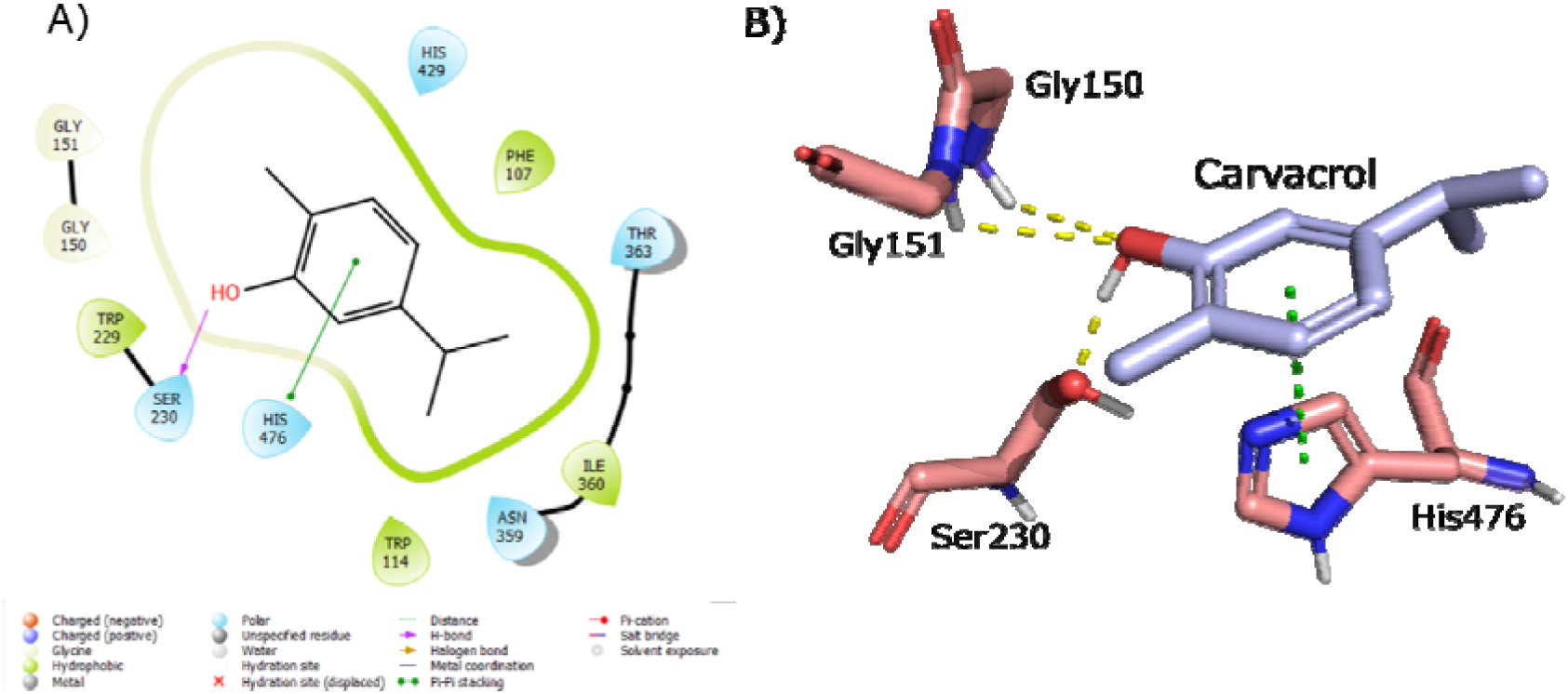
Binding sites of carvacrol on *R. microplus* acetylcholinesterase. 2D (A) and 3D (B) representation of ligand-protein interactions. In the 3D representation, residues and ligands are colored according to atom type (protein carbon, gray; oxygen, red; nitrogen, blue. Protein-ligand interactions are represented by dashed lines: hydrogen bridge interactions are colored in yellow, π→π interactions are colored in green—images were generated using PyMOL software.

We want to emphasize that the low acaricidal efficacy against the larval stage of ticks compared to the other 13 metabolites evaluated in this study can be explained by the structural characteristics observed in the exoskeleton under an optical microscope. The cuticle of the *Ornithodoros erraticus* larva is rougher and has a dense network, whereas in the fed adult stage of the tick, it induces stretching and increases flexibility and transcuticular absorption (Arrieta et al., 2013). Furthermore, arthropods possess an exoskeleton composed of esterified lipids and proteins, such as chitin (Beys-da-Silva et al., 2020). In the case of *R. microplus*, this could exert variable mechanical stress on the cuticle throughout its different stages. This structural difference suggests that i) the acaricidal efficacy of the metabolites is limited by an external physical barrier, determined by cuticular permeation kinetics (LogP and molecular weight), and ii) the permeation of compounds through the cuticle is easier in engorged adult ticks than in larvae. Consequently, these properties will determine the effective concentration, which would explain why metabolites with high molecular binding did not result in a significant percentage of larval mortality.

From a structural standpoint, carvacrol’s high efficacy is due to its lipophilic profile and low molecular weight. The presence of an aromatic ring that confers lipophilic properties, an OH group, and its relatively small size facilitate its diffusion through the lipid architecture of the tick’s exoskeleton, overcoming the external permeation barrier (Figure 10). A notable variability in lethal concentrations can be observed between the findings of this investigation (570 and 683 ppm) and other studies reported to date for this same metabolite. For example, Novato et al. (2015) reported substantially high LC_50_ values for *Amblyomma sculptum* (3490 ppm) and *Dermacentor nitens* (3330 ppm) (Novato et al., 2015). Other studies have reported greater sensitivity, with concentrations ranging from 830 ppm in *R. microplus* larvae (Navarro-Rocha et al., 2018), and a range of 1420 to 1760 ppm depending on the type of susceptible or resistant strain of *R. microplus* (Novato et al., 2022). This acaricidal activity positions carvacrol as a highly potent candidate and optimizes the cost-benefit ratio by requiring a smaller amount of active ingredient for the development of ixodicides.

**Figure 10.**
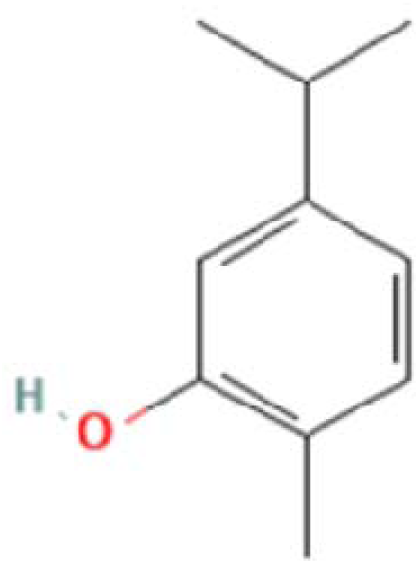
Chemical structure of carvacrol (A). Image obtained from PubChem.

The difference observed in the LC_50_ values of carvacrol between the two methods evaluated (LV and immersion) suggests that bioavailability and the surface area of exposure influence the compound’s effect. The larval immersion method allows for total cutaneous exposure, which increases transcuticular absorption, whereas in the LV method, uptake of the ixodicide is limited to contact with the tarsus of the tick’s legs during movement (Castro-Janer et al., 2009; Sabatini et al., 2001). Another relevant factor is the latency period in mortality readings; while conventional literature reports mortality at 24 h, our research found that a longer exposure period, up to 72 h, is essential to assess the full toxic effect of carvacrol. This sustained effect on mortality contrasts with the neurotoxic action of ethion, which exhibits a rapid acaricidal effect due to acute inhibition of AChE, but, in the long term, its lethality is negligible. In agreement with Duque et al. (2023), secondary metabolites exhibit prolonged toxicokinetics, possibly related to multiple mechanisms of action (Duque et al., 2023), such as the inhibition of mitochondrial electron transport chain observed in this study, which accumulate and affect the parasite’s homeostasis. Consequently, extensive studies on the acaricidal effect allow for more accurate LCLL estimates that are competitive with those of synthetic acaricides; furthermore, they demonstrate the true potential of these secondary metabolites as pest control agents.

Regarding the mechanism of action of carvacrol, as assessed by NADH oxidase and succinate oxidase assays, the results in Figure 6 indicate a preference for electron transport chain activity. The inability to detect an oxygen consumption response via complex II or succinate oxidase, even at high protein concentrations (2 mg/mL, *Figure S2* of the supplementary material), confirms that succinate oxidase has substantially lower activity compared to NADH oxidase. This finding suggests that mitochondrial metabolism in *R. microplus* larvae depends predominantly on NADH oxidation by complex I. This metabolic profile can be explained by the natural behavior of tick larvae. After hatching, the larvae climb onto vegetation and remain on the undersides of leaves, avoiding direct sunlight and entering a state of inactivity until they detect a host. During this resting period, the larvae can survive for up to eight months by minimizing their metabolic activity (Leal et al., 2018). Furthermore, when larvae are exposed to environmental conditions different from those of their natural habitat, such as refrigeration at 4 °C, they can enter a diapause-like state in which metabolic activity is further reduced (Leal et al., 2020).

In ticks, diapause is characterized by larval inactivity, a delay in the female’s feeding, a delay in metamorphosis, and a delay in oviposition or egg maturation. This state can be triggered by unfavorable environmental conditions, such as extreme temperatures, low humidity, host scarcity, or disruptions in the life cycle caused by external factors (Belozerov, 1982; Korotkov, 2016). During diapause, the marked reduction in physiological activity leads to a decrease in metabolic rate, and, as in hibernation, the activity of the mitochondrial enzyme complex is affected (Mathers et al., 2017). A similar metabolic suppression has been described in other organisms; studies by Duerr and Podrabsky (2010) demonstrated that embryos of the fish *Austrofundulus limnaeus* experience significant reductions in mitochondrial metabolism during diapause, including a decrease in the activity of complexes I, II, and IV, as well as ATP synthase (Duerr & Podrabsky, 2010).

The inhibitory effect of carvacrol on NADH oxidase suggests an interruption in electron flow along the mitochondrial electron transport chain, specifically at complexes I, III, and IV (Figure 6). During oxidative phosphorylation, carvacrol was shown to inhibit NADH oxidation coupled to ubiquinone reduction in complex I, which decreased the electron flux toward the artificial ferricyanide acceptor compared to the control. In contrast, carvacrol did not affect succinate dehydrogenase activity (Figure 6). This confirms that the metabolic capacity of complex II is intrinsically lower than that of complex I in this model. Now, when electrons were supplied via NADH (complex I) to assess the effect on ubiquinol-cytochrome c reductase, carvacrol reduced the enzyme’s activity by 30.0 ± 3.3%. Complex II (succinate dehydrogenase) was also used as an electron source in this enzyme to avoid bias, which had no effect on the pathway. In fact, when evaluating the activity of the succinate-cytochrome c reductase enzyme, carvacrol achieved an inhibition of the cytochrome c reduction process of 23.1 ± 4.6%, confirming that the decrease in activity is attributed exclusively to interference with complex III. As the final step of the electron transport chain, cytochrome c oxidase (Complex IV) showed significant sensitivity to the treatment, with an inhibition of 34.83 ± 9.61%. Unlike the transport complexes, the effect of carvacrol on ATP synthase did not produce statistically significant changes, demonstrating a selective toxicity profile that disrupts electron flow and causes the consequent loss of the proton gradient, leaving ATP synthesis intact, although this is limited by the proton gradient required due to the previous blockage.

This interference between complexes I and III can be explained by carvacrol’s structural affinity for ubiquinone binding sites, or by its ability to alter the lipid environment of transmembrane proteins due to its hydrophobic nature. The phenolic ring destabilizes lipid interactions and/or competes for ubiquinone binding sites, and its small size facilitates its interaction with cell membranes (both cytosolic and mitochondrial), thereby altering their fluidity and organization and destabilizing the catalytic functions of the enzyme complexes (Nelson et al., 2021). Furthermore, hydroxyl groups are active sites in many enzymatic reactions, as they can interact with various polar amino acid residues, which is important for the enzyme’s catalytic activity. Thus, they form hydrogen bonds with residues such as glycine, methionine, aspartic acid, and threonine; this can have serious consequences, such as altering the enzyme’s binding to its substrate (Hadjiivanov, 2014). Molecular docking predicted that carvacrol interacts with complexes I, III, and IV via hydrogen bonds between its OH group and the amino acids Gly249-Ile247 -Met250, Asp267-Trp209, and Thr315 of each complex, respectively, as well as a π-π interaction between the carvacrol ring and that of Trp209 (Figure 11). In this regard, the utility of the molecular findings obtained through molecular docking is supported by the efficacy observed in natural hosts, where the formulation used enhances and preserves the stability of the metabolite, promoting its permeability and interaction with pharmacological targets in the mitochondria (Anjos et al., 2024).

**Figure 11.**
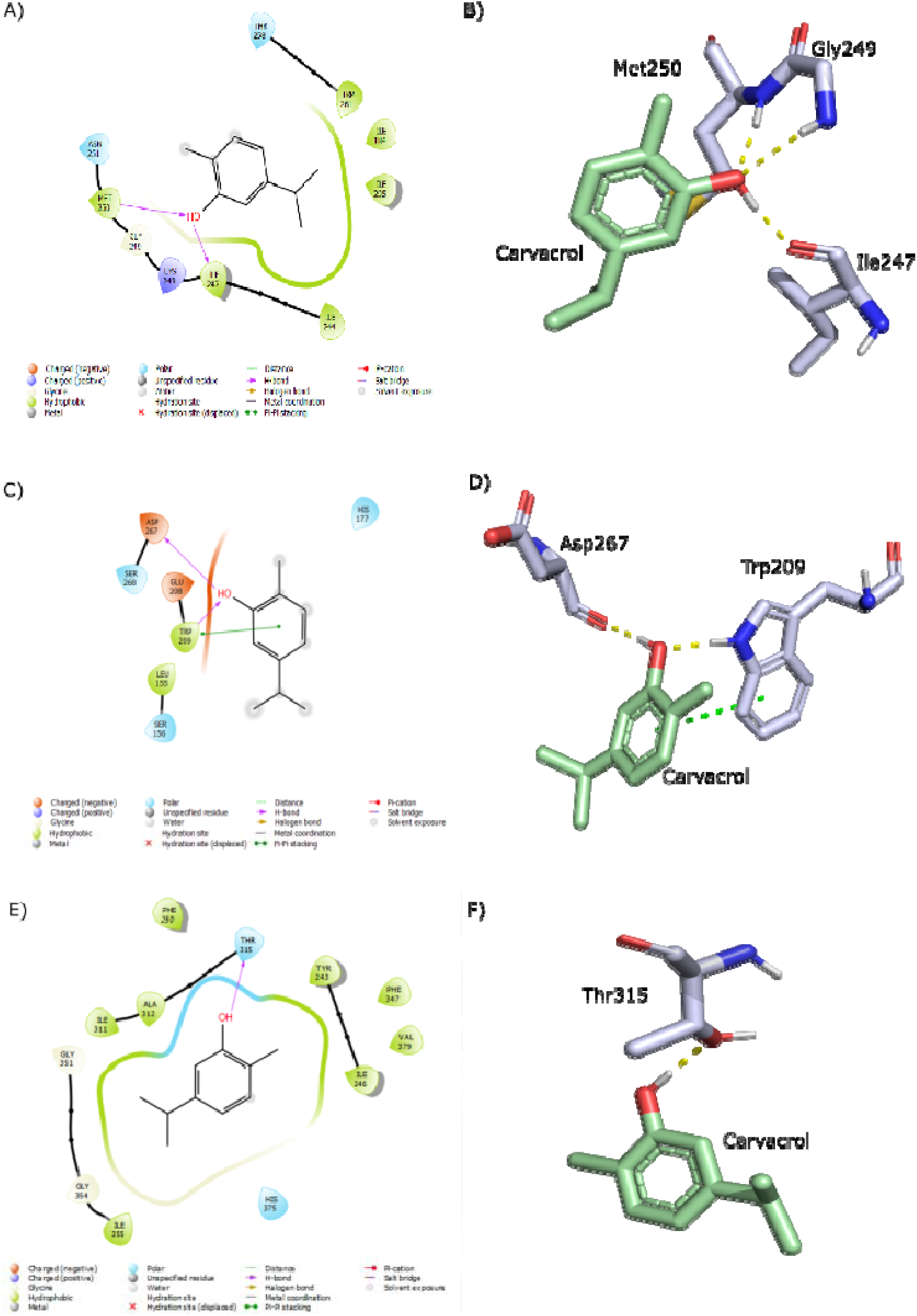
Binding sites of carvacrol (A, B) with NAD4 (chain 4 of complex I, ID: V9MMD6), (C, D) with mitochondrial Rieske (complex III, ID: A0A6M2CSX0), and (E, F) with COX1 (complex IV, ID: V9MLX4) in *R. microplus*. Residues and ligands are colored according to atom type (protein carbon, gray; oxygen, red; nitrogen, blue). Protein-ligand interactions are represented by dashed lines; hydrogen bond interactions are colored yellow, and π→π interactions are colored green. Images generated using PyMOL software.

The efficacy observed in the spray and pour-on formulations, compared to the group treated with ethion, is due to the presence of the carriers (ethanol and mineral oil) used. These promote absorption, temporarily disrupt the lipid architecture of the tick’s epicuticle, reduce surface tension, and/or facilitate homogeneous dispersion over the tick’s cuticle. Furthermore, the difference in efficacy between the spray formulation (20% every 10 days, p < 0.001) and the pour-on formulation (5% every 10 days, p = 0.093) suggests that the rate of absorption is key to the acaricidal effect. This difference is explained by the presence of ethanol, which serves as a vehicle and as a promoter of penetration into the cuticle; that is, it helps the compounds cross biological membranes. It has been documented that ethanol interacts with the lipid layers of the cuticle and alters its structure through interaction of the hydroxyl group, destabilizing the hydrogen bonds between its ceramides; thus, it alters the surface barrier of the exoskeleton and allows carvacrol to pass through the mite’s cuticle (Gamal et al., 2023). Another difference is that the pour-on application relies on the product’s diffusion through the lipids of the livestock’s dermis, whereas the spray technique achieves complete coverage of the host and, consequently, of the tick, maximizing the concentration of carvacrol and facilitating passive diffusion toward the mitochondrial and cholinergic targets identified through molecular docking. This reinforces the point mentioned earlier: the greater the direct exposure and surface area of contact, the better the activity of carvacrol. It also suggests pharmacological robustness dependent on the delivery vehicle and adequate saturation of physical barriers, such as the cuticle.

## 5. Conclusions and Future Perspectives

An integrated and systematic strategy was established for the identification of new acaricides through molecular docking and the use of AlphaFold models. Laboratory validation and preliminary field evaluation demonstrated that carvacrol exerts a dual mode of action: inhibition of complexes I, III, and IV of the mitochondrial electron transport chain and impairment of the central nervous system through acetylcholinesterase inhibition. The application of the ethanol formulation via spraying reduced the parasite load in the animals, surpassing the efficacy of the commercial product (Etion). As future prospects for this research, it is proposed to expand the field trial to understand the effect of carvacrol throughout the tick’s reproductive cycle and to conduct stability studies to evaluate the effect of temperature and radiation on carvacrol concentration under real-world conditions. Improve the formulation by implementing advanced delivery systems, for example, based on nanoformulations like nanoemulsions or invasomes, with the aim of enhancing penetration into the cuticle and the long-term stability of carvacrol as the active ingredient.

## Author contributions: CRediT

**Gloria S. Avendaño-Mora**: Conceptualization, Metodology, Formal analysis, Investigation, Writing - Original Draft, Writing - Review and Editing. **Andrés F. Cabezas-Cárdenas**: Metodology, Investigation, Writing - Original Draft. **Bethsy N. Alfonso-Núñez**: Metodology, Investigation, Writing - Original Draft. **Erika M. Celis Celis**: Metodology, Investigation, Writing - Review and Editing. **Cristhian Camilo Otero**: Investigation, Metodology. **Julián M. Botero Londoño**: Metodology, Resources, Writing - Review and Editing. **Wendy N. Triana**: Metodology, Investigation. **María Carolina Velásquez-Martínez**: Review and Editing. **Stelia C. Mendez-Sanchez**: Conceptualization, Metodology, Resources, Writing - Review and Editing, Supervision, Project administration, Funding acquisition. **Jonny E Duque**: Conceptualization, Metodology, Formal analysis, Resources, Writing - Review and Editing, Supervision, Project administration, Funding acquisition.

## Acknowledgments

Special thanks are extended to Dr. Luis Carlos Vesga Gamboa for his training in the use of Schrödinger Maestro software and molecular docking approaches. The authors also thank Gustavo Adolfo Rincón Sandoval for his technical support at the Medical Entomology Laboratory of CINTROP, and to Karen Yineth Prada Campos for her valuable collaboration in the preparation of the figures of the acaricidal methodologies against larvae.

## Appendix A. Supporting information

