## Supplemental material for "Carvacrol as a dual-target acaricidal candidate against the cattle tick *Rhipicephalus (Boophilus) microplus*: molecular docking, mechanistic validation, and early field testing"

**Supplementary material**

**Tables section**

**Table S1.** Coordinates of the box thrown by the site map for each subunit.

| **Subunit** | **X Coordinate** | **Y Coordinate** | **Z Coordinate** |
| --- | --- | --- | --- |
| NAD4 subunit, Chain 4 of complex I | 219 | 161 | 146 |
| Iron-sulfur subunit, mitochondrial | -9 | -17 | -47 |
| Mitochondrial Rieske subunit | 33 | 34 | 37 |
| COX1 subunit Subunit 1 | 10 | 3 | 7 |
| AChE | 10 | 75 | 47 |

**Table S2**. Please see the attached Excel file named “List of metabolites identified from the plant species reported by Benelli et al.”

**Table S3**: Pre-selected multitarget secondary metabolites as potential inhibitors of mitochondrial CTE and AChE protein subunits in R. microplus ticks.

| **Mitochondrial ETC complex and AChE** | **Protein subunit or enzyme** | **MS with possible inhibitory activity** | **Docking score** | **Affinity energy** |
| --- | --- | --- | --- | --- |
| I | NADH-ubiquinone oxidoreductase | Harman | -34.050 | -4.42 |
|  |  | Quercetin | -29.090 | -4.56 |
|  |  | 3,4-dihydroxyphenylacetic acid | -21.450 | -4.20 |
|  |  | Agnusida | -9,96 | -59,29 |
|  |  | Clorogenic acid | -8,76 | -32,96 |
|  |  | Catechin | -7,82 | -38,86 |
|  |  | Hesperidin | -7.457 | -78.66 |
|  |  | Nepitrine | -7,40 | -57,29 |
|  |  | Aucubin | -7,08 | -36,56 |
|  |  | Andrographolide | -6.915 | -67.00 |
|  |  | Rosmarinic acid | -6,82 | -49,16 |
|  |  | Caffeic acid | -6,58 | -8,89 |
|  |  | Hydrobenzoic acid | -6.091 | -46.68 |
|  |  | Vitamin A | -5.382 | -62.31 |
|  |  | Epicatechin | -5.239 | -44.20 |
|  |  | H-fluorene-4-carboxylic acid | -5.215 | -42.79 |
|  |  | Cuminol | -5.191 | -45.41 |
|  |  | Valenceno | -5.177 | -43.19 |
|  |  | Gallic acid | -4.597 | -25.52 |
| II | Succinate dehydrogenase[ubiquinone] Fe-S | Nepitrine | -8,92 | -69,85 |
|  |  | Epicatechin | -8.136 | -54.93 |
|  |  | Rosmarinic acid | -7,08 | -53,13 |
|  |  | Quercetin | -6.683 | -45.06 |
|  |  | Eriodictol | -6.380 | -47.40 |
|  |  | Hesperidin | -6.327 | -47.42 |
|  |  | Catechin | -6,06 | -45,15 |
|  |  | Atropine | -5,69 | -44,76 |
|  |  | Gallic acid | -5.397 | -43.95 |
|  |  | Syringaldehyde | -5.330 | -42.98 |
|  |  | 1-phenyl-1,2-propanedione | -5.293 | -40.89 |
|  |  | Dihydropiplartine | -5,14 | -41,87 |
|  |  | Naringenin | -5.134 | -44.98 |
| III | Cytochrome b-c1 | Chlorogenic acid | -6.127 | -41.42 |
|  |  | Naringenin | -6.045 | -35.81 |
|  |  | Pinocembrin | -5,76 | -32,32 |
|  |  | Hesperidin | -4.971 | -32.81 |
|  |  | Quercetin | -4.837 | -35.81 |
|  |  | Luteolin | -4,51 | -32,54 |
|  |  | Hydrobenzoin | -4.432 | -29.06 |
|  |  | Patuletine | -4,41 | -40,58 |
|  |  | Eriodictol | -4.223 | -31.31 |
|  |  | Carvacrol | -4.180 | -28.74 |
|  |  | Epicatechin | -4.099 | -33.24 |
|  |  | Aucubin | -3,89 | -27,27 |
|  |  | Gallic acid | -3.888 | -21.74 |
| IV | Cytochrome C oxidase | H-fluorene-4-carboxylic acid | -8.136 | -43.94 |
|  |  | 3,4-Dihydroxyphenylacetic acid | -7.602 | -36.63 |
|  |  | Quinic acid | -7.372 | -35.99 |
|  |  | Gallic acid | -7,20 | -30,82 |
|  |  | Trans-ferulic acid | -6.712 | -52.17 |
|  |  | Crisina | -6.666 | -54.28 |
|  |  | Flavanone | -6.559 | -43.93 |
|  |  | 3-Hydroxyflavone | -6.545 | -46.49 |
|  |  | Citric acid | -6.328 | -2.36 |
|  |  | Imperatorin | -6.305 | -60.78 |
|  |  | 2,5-dihydroxybenzoic acid | -6.079 | -39.91 |
|  |  | Phthalic acid | -5.925 | -19.06 |
|  |  | 1-phenyl-1,2-propanedione | -5.388 | -31.66 |
|  |  | Flavone | -5.362 | -53.45 |
|  |  | Hydrobenzoin | -5.157 | -54.84 |
|  |  | 3',5'-Dimethoxyacetophenone | -5.038 | -34.77 |
|  |  | Caffeic acid | -5.010 | -27.67 |
|  |  | D-Limonene | -4.504 | -21.27 |
|  |  | Linalool | -4.398 | -30.16 |
|  |  | Eugenol | -4.356 | -29.16 |
|  |  | Gamma-terpinene | -4.146 | -17.91 |
|  |  | Geraniol | -3.770 | -25.72 |
|  |  | Carvacrol | -3.749 | -28.45 |
|  |  | Citral | -3.703 | -28.96 |
|  |  | Stylbene | -3,70 | -30,66 |
| AChE | Acetylcholinesterase | Hesperidin | -9.524 | -44.97 |
|  |  | Aucubin | -8,05 | -31,96 |
|  |  | Quercetin | -6.559 | -32.45 |
|  |  | Epicatechin | -6.460 | -26.84 |
|  |  | Agnuside | -6,23 | -47,84 |
|  |  | Andrographolide | -5.797 | -47.52 |
|  |  | Luteolin | -5,49 | -28,63 |
|  |  | Eriodictol | -5.457 | -30.88 |
|  |  | Junipegenin A | -5,43 | -35,47 |
|  |  | Quinic acid | -5.290 | 3.25 |
|  |  | Naringenin | -5.161 | -24.91 |
|  |  | Catechin | -4,90 | -29,85 |
|  |  | Gallic acid | -4.341 | -7.41 |
|  |  | Alolicoisoflavone A | -4,69 | -37,53 |

**Table S4.** Summary of Oxford Nanopore sequencing results showing the total bases and reads obtained for each barcode.

| **Sample** | **Total Bases**  **(Mb)** | **Passed Bases (%)** | **Total Reads**  **(k)** | **Passed Reads (%)** |
| --- | --- | --- | --- | --- |
| S7374 | 31.88 | 58.8 | 75.33 | 60.8 |
| S7576 | 23.86 | 47.4 | 49.25 | 51.4 |
| S7376 | 36.51 | 51.2 | 91.39 | 57.1 |

Summary of Oxford Nanopore sequencing results showing the total bases and reads obtained for each sample. S7374: COX-1 amplicon of *Rhipicephalus microplus* generated with primers F73/R74; S7576: COX-1 amplicon of *Rhipicephalus microplus* generated with primers F75/R76; S7376: COX-1 amplicon of *Rhipicephalus microplus* generated with primers F73/R76.

**Table S5.** Please see the attached Excel file named “*Raw_Data*”. Corresponding results of the acaricidal tests for the 13 metabolites with activity resulting in less than 20% mortality.

**Table S6.** Summary of negative binomial GLMM results for field test

| **Predictor** | **Estimate** | **SE** | **z** | **p-value** |
| --- | --- | --- | --- | --- |
| (Intercept) | 4.9816 | 0.1135 | 43.889 | <0.001 |
| Treatment: Pour-on formulation | -0.9742 | 0.0885 | -11.006 | <0.001 |
| Treatment: Spray vehicle | -0.8316 | 0.1851 | -4.492 | <0.001 |
| Treatment: Pour-on vehicle | -0.3971 | 0.0797 | 4.985 | <0001 |
| Treatment: Ethion | -0.6704 | 0.1939 | -3.458 | <0.001 |
| Day10 | -0.2234 | 0.0285 | -7.855 | <0.001 |
| Sex (Male) | 0.5088 | 0.1818 | 2.799 | 0.005 |
| Age_c | -0.0361 | 0.0277 | -1.302 | 0.193 |
| Pour-on formulation ˟ Day10 | 0.1723 | 0.0417 | 4.136 | <0.001 |
| Spray vehicle ˟ Day10 | 0.3639 | 0.0414 | 8.790 | <0.001 |
| Pour-on vehicle ˟ Day10 | 0.4101 | 0.0459 | 8.940 | <0.001 |
| Ethion ˟ Day10 | 0.3372 | 0.0417 | 8.090 | <0.001 |

Results of the negative binomial generalized linear mixed model (log link) evaluating the effects of treatment, time, and their interaction on tick counts. Time was modeled in 10-day units. Treatment levels included spray formulation, pour-on formulation, spray vehicle, pour-on vehicle, and ethion. Animal ID was included as a random intercept. Interaction terms (Treatment × Day10) represent treatment-specific temporal slopes relative to the reference category. SE, standard error.

**Table S7.** The estimated rate of change in tick counts over time under each treatment

| **Treatment** | **Slope (log scale)** | **% change per-10 days** | **95% CI (%)** | **p-value** |
| --- | --- | --- | --- | --- |
| Spray formulation | -0.223 | -20.0 | (-24.4, -15.4) | < 0.001 |
| Pour-on formulation | -0.051 | -5.0 | (-10.5, 0.9) | 0.093 |
| Spray vehicle | 0.140 | +15.1 | (8.5, 22.1) | < 0.001 |
| Pour-on vehicle | 0.187 | +20.5 | (12.3, 29.3) | < 0.001 |
| Ethion | 0.114 | +12.0 | (5.6, 18.9) | < 0.001 |

Slopes were estimated from a negative binomial generalized linear mixed model. Percentage change was calculated as (exp(slope) − 1) × 100. Negative values indicate a reduction in tick counts over time. Time was scaled in 10-day units. CI, confidence interval.

**Table S8.** Pairwise comparisons of treatment-specific temporal slopes derived from the negative binomial generalized linear mixed model

| **Contrast** | **Estimate** | **SE** | **df** | **LCL** | **UCL** | **z-ratio** | **p-value** |
| --- | --- | --- | --- | --- | --- | --- | --- |
| Spray formulation – Pour on formulation | -0.1723 | 0.0417 | Inf. | -0.2859 | -0.0587 | -4.136 | 0.0003 |
| Spray formulation – Spray vehicle | -0.3639 | 0.0414 | Inf. | -0.4769 | -0.2510 | -8.790 | <0.0001 |
| Spray formulation – Pour on vehicle | -0.4101 | 0.0459 | Inf. | -0.5352 | -0.2850 | -8.940 | <0.0001 |
| Spray formulation – Ethion | -0.3372 | 0.0417 | Inf. | -0.4509 | -0.2235 | -8.090 | <0.0001 |
| Pour on formulation – Spray vehicle | -0.1916 | 0.0428 | Inf. | -0.3084 | -0.0749 | -4.478 | 0.0001 |
| Pour on formulation – Pour on vehicle | -0.2378 | 0.0471 | Inf. | -0.3664 | -0.1092 | -5.044 | <0.0001 |
| Pour on formulation – Ethion | -0.1649 | 0.0431 | Inf. | -0.2823 | -0.0474 | -3.829 | 0.0012 |
| Spray vehicle – Pour on vehicle | -0.0462 | 0.0469 | Inf. | -0.1741 | 0.0818 | -0.984 | 0.8626 |
| Spray vehicle – Ethion | 0.0267 | 0.0428 | Inf. | -0.0900 | 0.1435 | 0.625 | 0.9712 |
| Pour on vehicle – Ethion | 0.0729 | 0.0471 | Inf. | -0.0557 | 0.2015 | 1.546 | 0.5323 |

Contrasts were Tukey-adjusted for multiple comparisons. Estimates represent differences in log-scale slopes between treatment groups. SE, standard error; df, degrees of freedom; LCL, lower confidence limit; UCL, upper confidence limit.

**Figures section**

**Figure S1**. Agarose gel electrophoresis of PCR amplicons obtained.

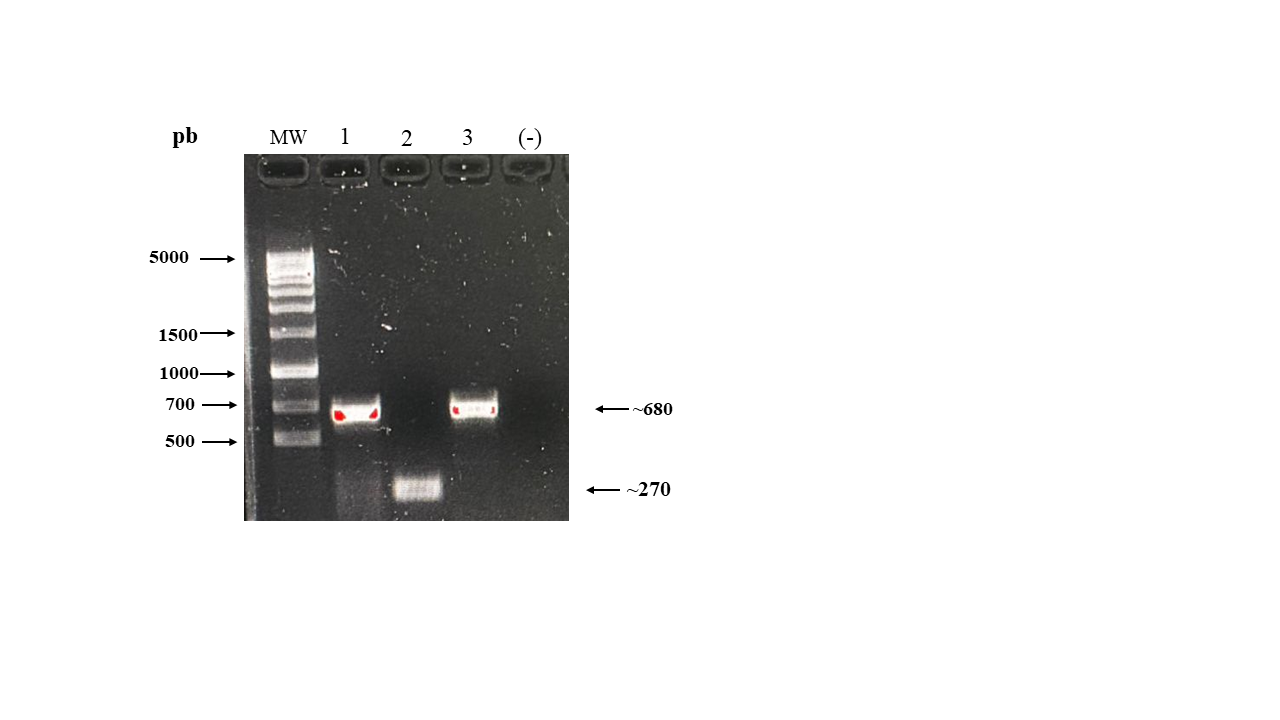

MW: molecular weight marker. Lane 1: PCR amplification of *Rhipicephalus microplus* DNA using primers F73 and R74 (S7374). Lane 2: PCR amplification of *Rhipicephalus microplus* DNA using primers F75 and R76 (S7576). Lane 3: PCR amplification of *Rhipicephalus microplus* DNA using primers F73 and R76 (S7376). Lane (-): negative control.

**Figure S2.** Multiple sequence alignment of the Nanopore-derived consensus sequence (S7374) against reference sequences identified by BLASTn in Geneious Prime.

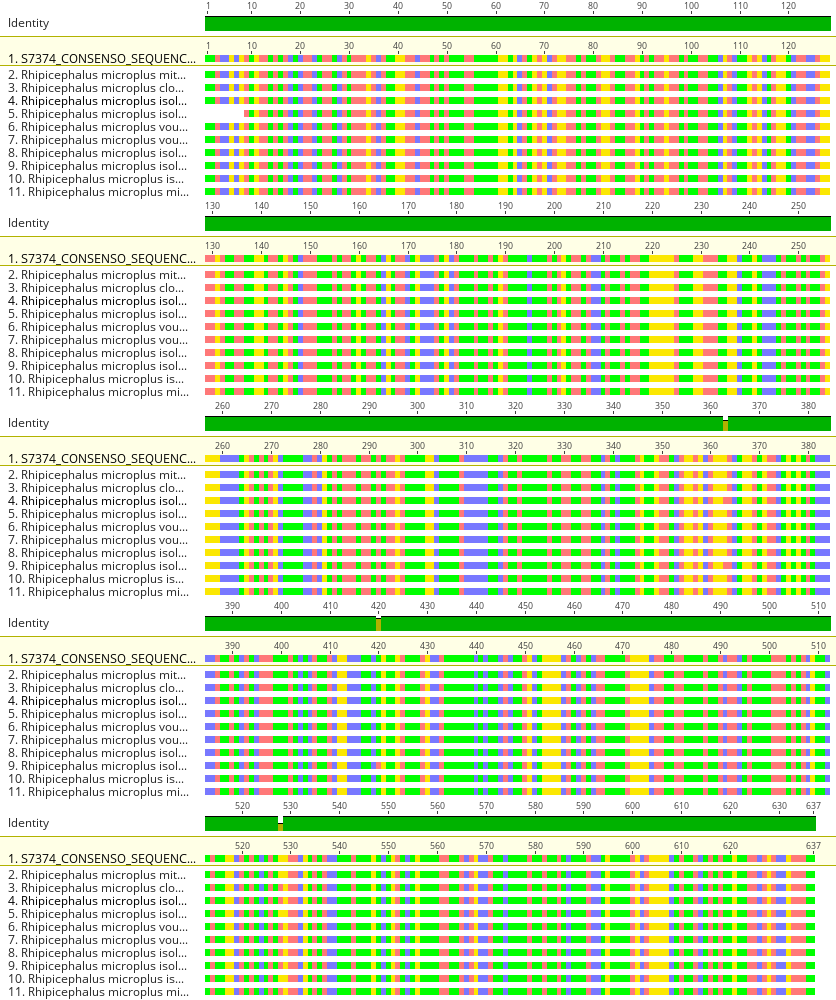

**Figure S3.** Multiple sequence alignment of the Nanopore-derived consensus sequence (S7576) against reference sequences identified by BLASTn in Geneious Prime.

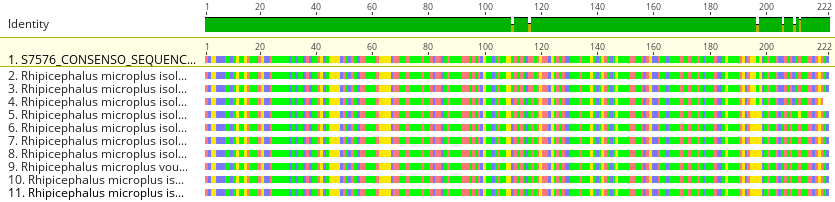

**Figure S4.** Multiple sequence alignment of the Nanopore-derived consensus sequence (S7376) against reference sequences identified by BLASTn in Geneious Prime

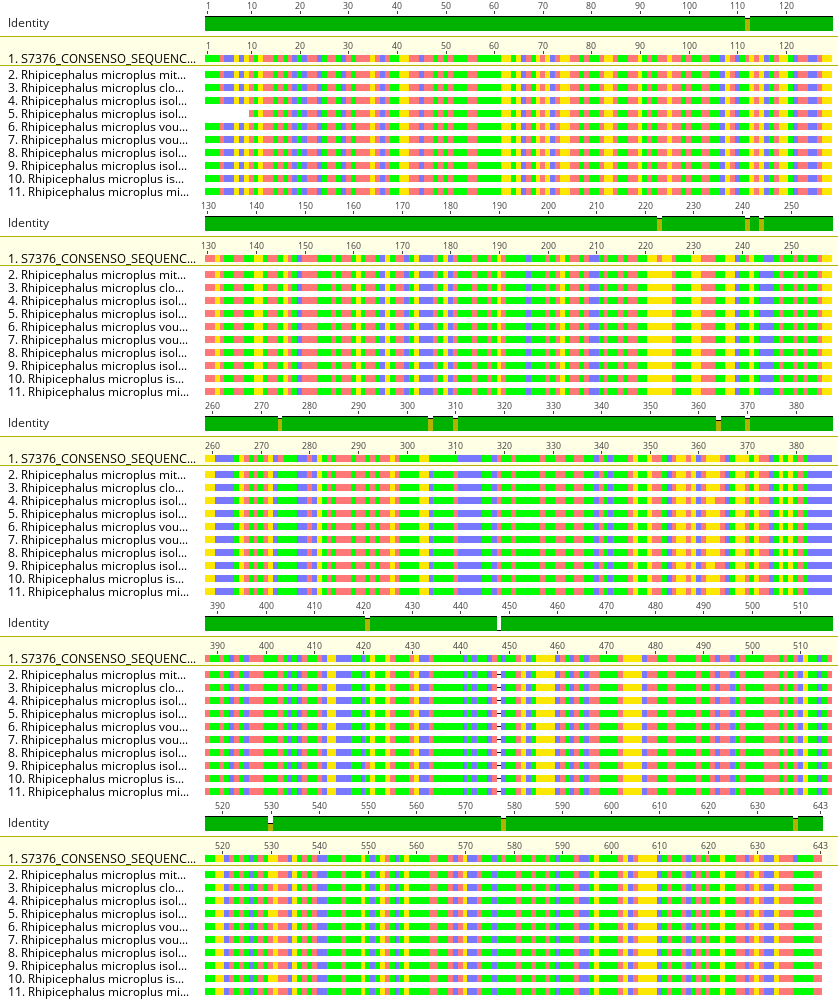

**Figure S5**. Lethality caused by carvacrol at diagnostic concentrations. Different letters (a, b, c, d) indicate significant differences from the negative control (p ≤ 0.05)

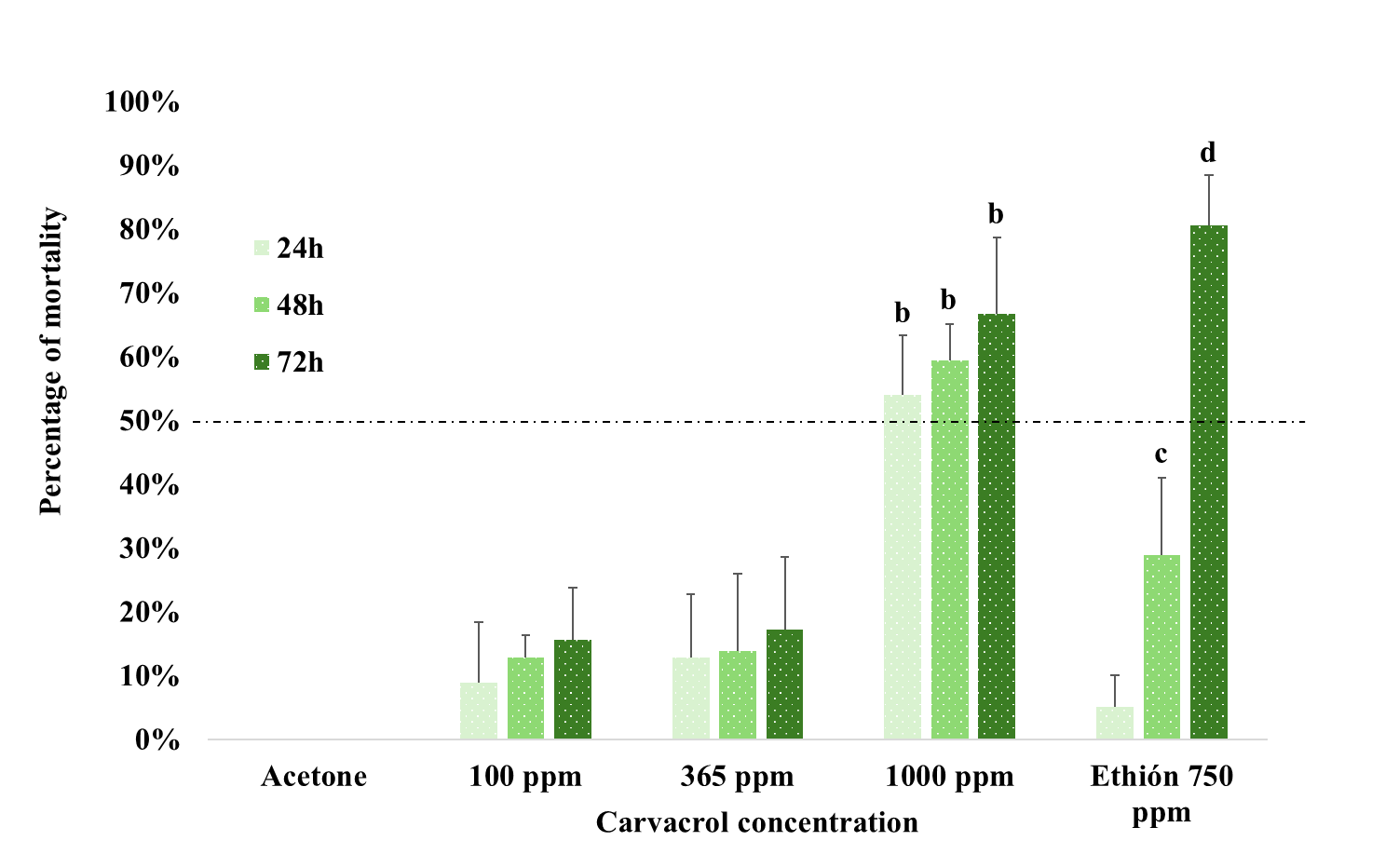

**Figure S6.** Succinate oxidase activity at different concentrations of total protein.

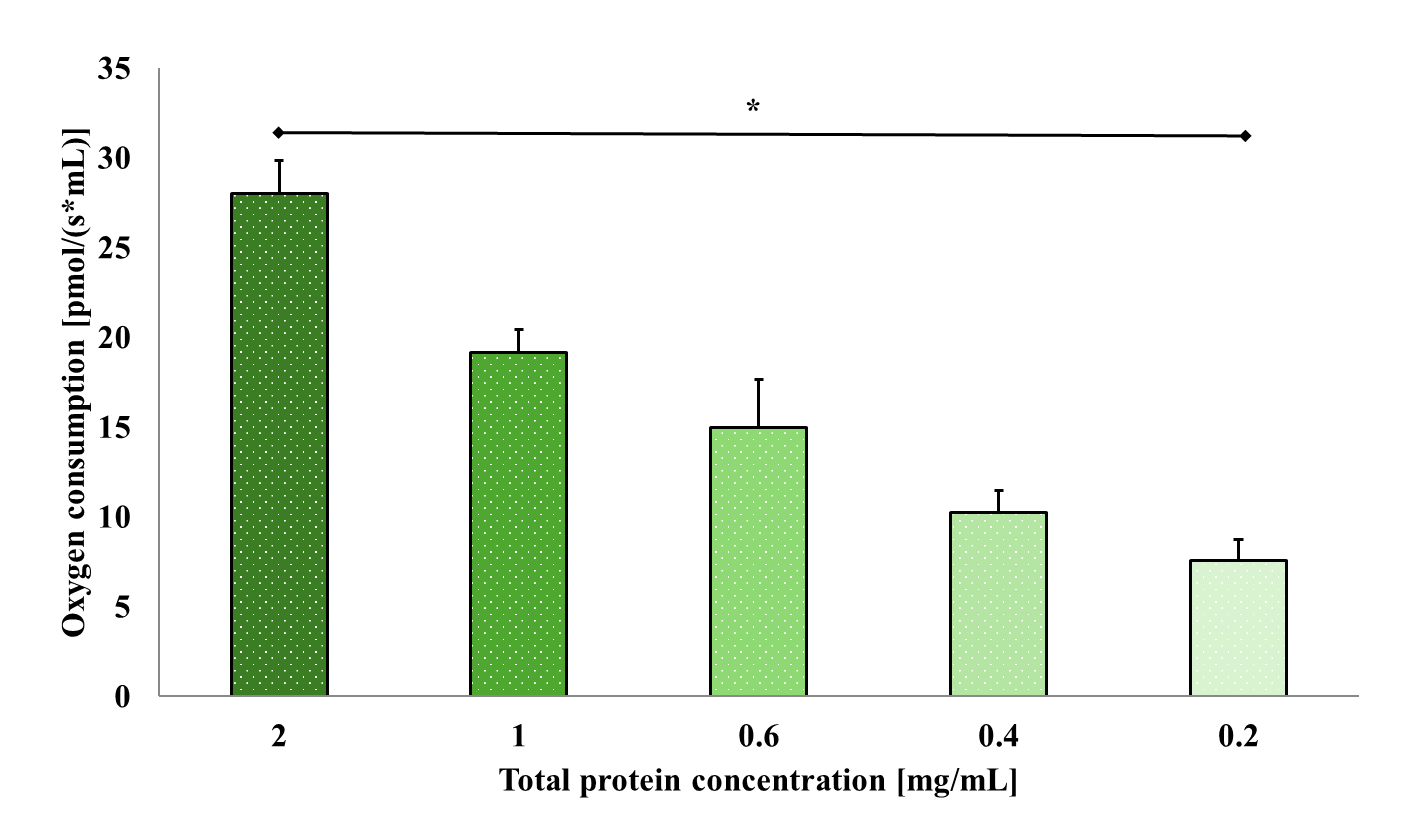

The same total protein concentration (0.6 mg/mL) was used for succinate oxidase assays; however, the activity was markedly lower. To evaluate the experimental feasibility of this enzyme, succinate oxidase activity was analyzed in the absence of carvacrol and at increasing total protein concentrations. As shown in Figure 6, the oxygen consumption rate increased proportionally with total protein concentration. At the maximum evaluated concentration (2 mg/mL total protein), the oxygen consumption rate reached 22.40 ± 1.03 pmol/(s·mL).

The Shapiro-Wilk normality test was performed (p > 0.05), followed by an ANOVA with Tukey’s multiple comparisons test. (*) indicates significant differences (p < 0.05). Statistically significant differences were observed across all total protein concentrations (p ≤ 0.05*).

**Figure S7.** Predicted tick counts over time by treatment.

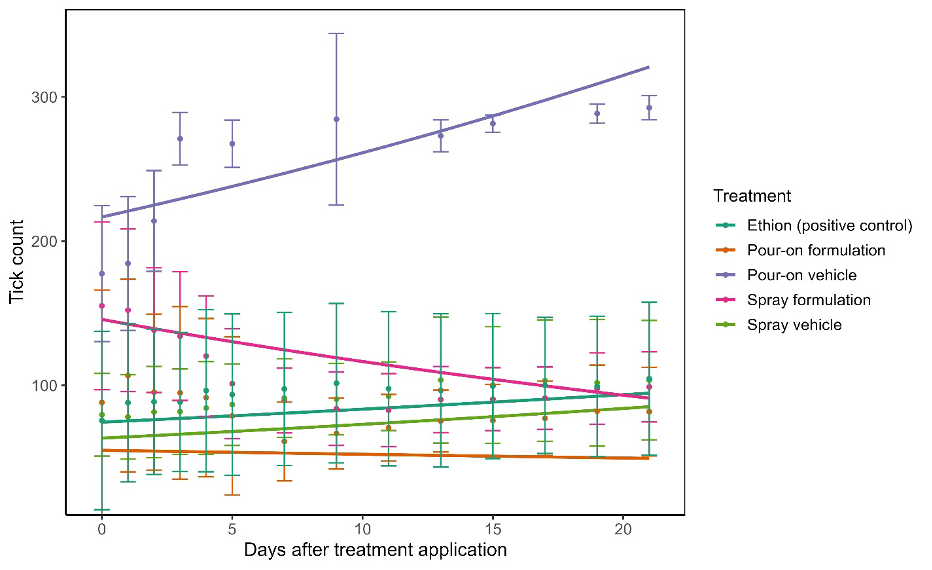

**Note:** Solid lines represent predicted values from the negative binomial generalized linear mixed model for each treatment. Points indicate observed mean tick counts at each sampling day, with error bars representing ± standard deviation.
